# PINK1 loss in astrocytes triggers inflammatory dysfunction and neuronal death

**DOI:** 10.64898/2026.06.04.729996

**Authors:** Gabriella Fiorino, Damian N. Di Florio, Deshaun H. Hadley, Owen A. Ross, Fabienne C. Fiesel, Wolfdieter Springer

## Abstract

Genetic loss of the mitochondrial control enzyme PINK1 leads to Parkinson’s disease, characterized by dopaminergic neuron degeneration and neuroinflammation, yet its role in glia remains poorly understood. To address this gap, we investigated how the function of astrocytes and their ability to support neurons is influenced by PINK1 deficiency. For the first time, we demonstrate that human astrocytes exhibit robust PINK1 activity. Next, the first bulk transcriptomic study of human PINK1 mutant astrocytes was performed followed by biochemical validation at the protein level, uncovering homeostatic collapse. Co-culture experiments demonstrated that this astrocyte dysfunction drives neuronal damage through non-cell-autonomous mechanisms. Notably, pharmacological enhancement of autophagy successfully mitigated this inflammatory secretome, indicating that mitochondrial quality control deficits are reversible. These findings establish an unexpected role for PINK1 in glial biology, reveal that astrocytes are vulnerable to mitophagy deficits, and highlight a novel mechanistic link connecting mitochondrial dysfunction, neuroinflammation, and neurodegeneration.

## INTRODUCTION

Parkinson’s disease (PD) is the second most common neurodegenerative disorder and among the fastest-growing neurological conditions globally*^(1)^*. While typically associated with aging, approximately 5-15% of cases are classified as early-onset Parkinson’s disease (EOPD), defined by clinical onset before 50 years of age*^(2)^*. Loss-of-function (LOF) mutations in the genes encoding the mitochondrial quality control enzymes PINK1 and PRKN are the most common causes of EOPD*^(2)^*. The ubiquitin kinase PINK1 and the E3 ubiquitin ligase PRKN/Parkin work together through a stress activated pathway to selectively label damaged mitochondria by phosphorylating serine 65 on ubiquitin (p-S65-Ub), tagging them for degradation via the autophagy-lysosome system (mitophagy)*^(3)^*. Nigrostriatal dopaminergic neurons are selectively vulnerable to the mitochondrial dysfunction induced by PINK1 or PRKN LOF due to their high metabolic demand, consistent pacemaking activity, unusually large axonal arborization, and particularly high levels of PINK1 and PRKN expression and activity*^(4–6)^*. However, the exact mechanisms causal of neuronal death and whether neurons constitute the only vulnerable central nervous system (CNS) cell type during PD remain unclear.

Besides dopaminergic neuron death, neuroinflammation is another hallmark of PD*^(7)^*. Astrocytic and microglial activation along with infiltration of CD4+ and CD8+ T cells are consistently observed at heightened levels in the substantia nigra pars compacta and striatum of PD patients, but not in unaffected brain regions*^(8, 9)^*. Traditionally, neuroinflammation is believed to be the consequence of glial response to dead or dying neurons. However, a growing body of literature supports the idea that glia may in fact drive neurodegeneration and that non-cell autonomous events may contribute to PD*^(10–14)^*. This is exemplified through recent work showing that microglial neurotoxicity can be triggered by another causative gene for PD, *LRRK2*, and that a *LRRK2* non-coding variant acts via microglia specific regulation of gene expression*^(10, 11)^*. Furthermore, female mice harboring a causal PD mutation in *LRRK2* exhibited macrophage exhaustion, which was linked to mitochondrial dysfunction and differential expression of PINK1 and PRKN via transcriptomic analysis*^(12)^*. Although mitochondrial dysfunction and inflammatory neurotoxicity are closely linked*^(15, 16)^*, little is known regarding the importance of the PINK1-PRKN pathway in non-neuronal CNS cell types.

PINK1 and PRKN are ubiquitously expressed across various CNS cell types to varying degrees*^(5, 17-19)^*, suggesting that they likely play an important role in the health and function of cell types beyond nigrostriatal dopaminergic neurons. Interestingly, one study reported a lack of gliosis in *PRKN* mutation carrier post-mortem human midbrain*^(20)^* while another identified a reduction of astrocyte reactivity markers in *PRKN* mutation carriers compared to healthy controls*^(21)^*. In contrast, the levels of these reactivity markers were heightened in idiopathic PD patient brain*^(21)^*. These results indicate that the PINK1-PRKN pathway is likely important for immune signaling regulation and mediating astrocyte reactivity. Astrocytes provide critical neuroprotection and have high bioenergetic demand, with mitochondrial densities in their cell bodies similar to that of neurons*^(22)^*. Therefore, lack of mitochondrial quality control may also induce astrocyte vulnerability.

In this study, using two independent gene-edited sets of PINK1 LOF mutants and their respective isogenic iPSCs, we experimentally defined transcriptional and functional alterations resulting from loss of PINK1 activity in astrocytes at physiologically relevant baseline conditions (i.e. without exogenous stressors). Loss of PINK1 function triggered release of neurotoxic and pro-inflammatory danger signals, deficits in the astrocytic inflammatory defense system, impaired proliferative capacity, astrocyte-neuron communication defects, and metabolic failure indicating homeostatic collapse and exhaustion. Pathway analysis further revealed dysfunction to mTOR signaling in PINK1 mutant astrocytes, which was subsequently functionally and biochemically validated. Of note, pharmacologically boosting autophagy activation via mTOR inhibition rescued the heightened astrocytic inflammatory signaling. Finally, co-culture systems via transwell insert revealed reduced viability in neurons cultured with PINK1 mutant astrocytes. These results indicate that PINK1 deficiency triggers widespread astrocyte dysfunction, thereby inducing neuronal damage and death in a non-cell autonomous manner.

## RESULTS

### Differentiation of PINK1 mutant iPSCs into astrocytes

To determine how PINK1 deficiency impacts human astrocyte function and the cells’ inherent ability to support neurons, two independent isogenic sets of PINK1 mutant iPSCs were differentiated into neural progenitor cells (NPCs) and then into astrocytes (iAS) using optimized protocols*^(23)^* (**Fig. 1A**). Set 1 consisted of a gene-edited PINK1 knockout (KO) line derived from its isogenic wild-type (isoWT) control. Set 2 consisted of patient-derived PINK1 I368N iPSCs and its gene-corrected, isogenic WT counterpart (isoWT), in which enzymatic activity was restored. We previously characterized the PINK1 I368N as a kinase dead mutation resulting from a deformed ATP-binding pocket and failure to properly stabilize full-length PINK1 protein*^(6, 24)^*. Pluripotency of each iPSC line was confirmed using the markers Nanog and Oct4 (**Fig. S1A**). The iPSCs were then converted into NPCs, resulting in ∼100% positivity for the NPC markers Pax6 and Nestin (**Fig. S1, B to D**). This suggests that loss of PINK1 gene expression or kinase activity does not impair NPC differentiation.

**Fig. 1.**
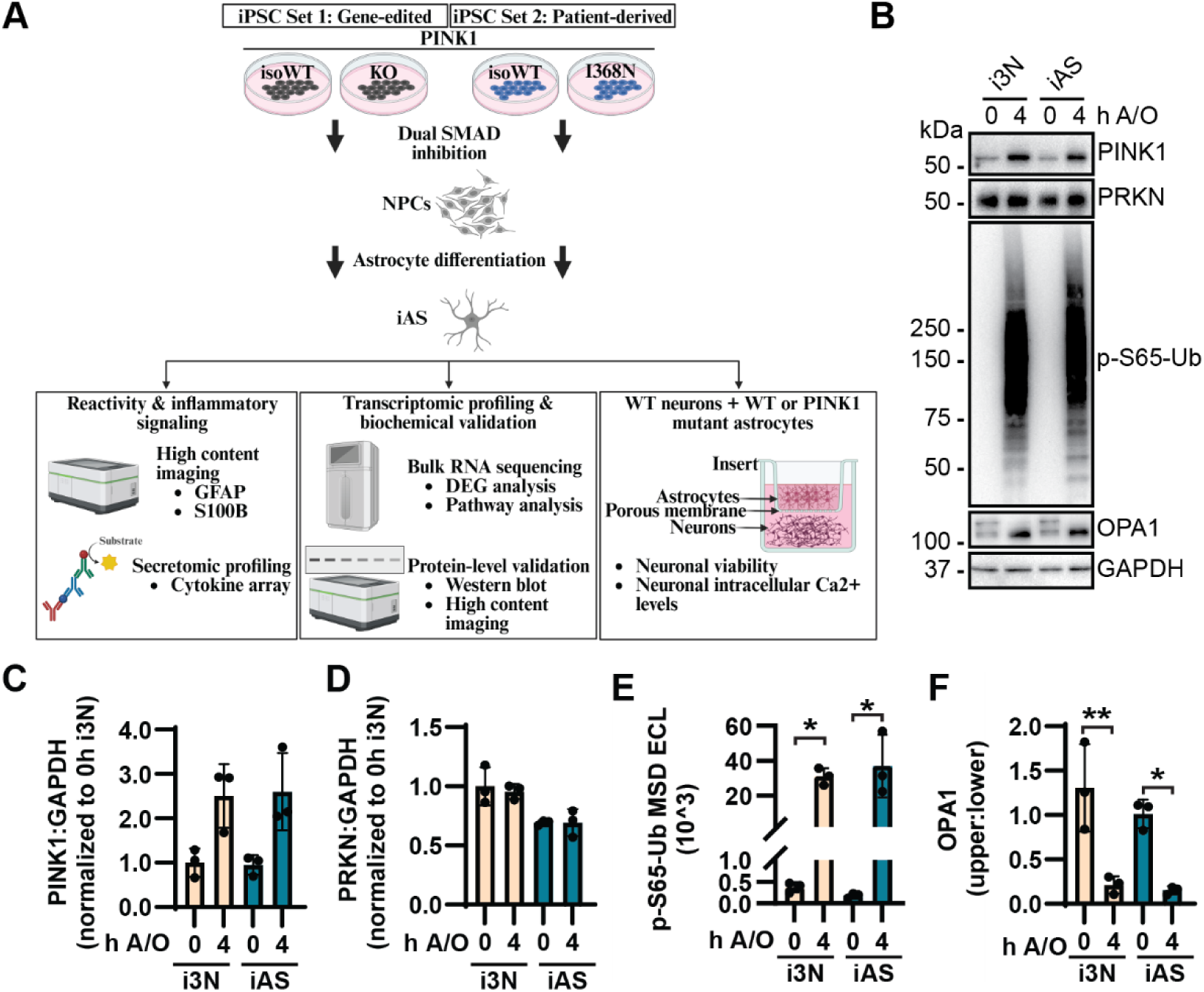
Astrocytes display robust levels of PINK1-PRKN signaling. **(A)** Schematic overview of the experimental workflow. NPC = neural progenitor cells. iAS = iPSC-derived astrocytes. **(B)** Representative western blot shows levels of PINK1, PRKN, p-S65-Ub, and OPA1 at baseline and in response to a combination of 4 µM antimycin and 10 µM oligomycin (A/O) treatment. **(C-D)** PINK1 and PRKN data points are displayed as each protein divided by GAPDH. **(E)** p-S65-Ub levels measured using MSD ELISA. **(F)** OPA1 levels were quantified as upper band divided by lower band. PINK1:GAPDH and PRKN:GAPDH data is represented as normalized to 0h i3N set to 1, while MSD results are displayed as background corrected raw data. The mean of three independent experiments for each cell type ± SD is depicted. Statistical analysis was performed using Two-Way ANOVA with Tukey’s multiple comparison test (**p<0.01, *p<0.05). The schematic was generated using BioRender.

The cells were subsequently differentiated into astrocytes to first define the level of PINK1-PRKN pathway activation in comparison to cortical-like i^3^ Neurons (i3N), as neurons are known to have high levels of mitophagy activation*^(5, 19, 25)^*. WT i3N and iAS were treated with antimycin and oligomycin (A/O) to activate PINK1 and PRKN to selectively label depolarized mitochondria for degradation. Interestingly, the levels of PINK1, PRKN, and the mitophagy tag p-S65-Ub were similar across both cell types (**Fig. 1, B to E**). Mitochondrial membrane depolarization was equally effective in response to A/O treatment as determined via cleavage of the long isoform of OPA1 (**Fig. 1F**). These findings indicate robust levels of PINK1-PRKN signaling in human astrocytes and are consistent with prior reports showing high levels of p-S65-Ub in rodent cortical astrocytes compared to other CNS cell types*^(26, 27)^*.

Next, the absence of PINK1 activity in both sets of mutant iAS was confirmed by measuring the levels of PINK1, PRKN, and p-S65-Ub under basal conditions and in response to mitochondrial stress (**Fig. 2A**). Effective mitochondrial membrane depolarization across genotypes was confirmed via OPA1 levels. As expected, the immunoblot results demonstrated a significant reduction in PINK1 levels from the PINK1 KO and patient-derived iAS compared to their respective isoWT (**Fig. 2B**). The absence of functional PINK1 was accompanied by a 3-4-fold accumulation of PRKN protein levels in both PINK1 mutants compared to their corresponding isoWT controls (**Fig. 2C**). These results are in agreement with our prior findings that PINK1 is critical for the activation and subsequent turnover of the E3 ubiquitin ligase PRKN*^(6)^*. PINK1-PRKN pathway activity was further quantified via a previously developed sensitive sandwich ELISA on a Meso Scale Discovery (MSD) platform for p-S65-Ub*^(28)^* (**Fig. 2D**). Both the PINK1 KO and I368N iAS displayed a complete lack of p-S65-Ub signal, providing further evidence of the absence of PINK1 kinase activity in both mutant sets.

**Fig. 2.**
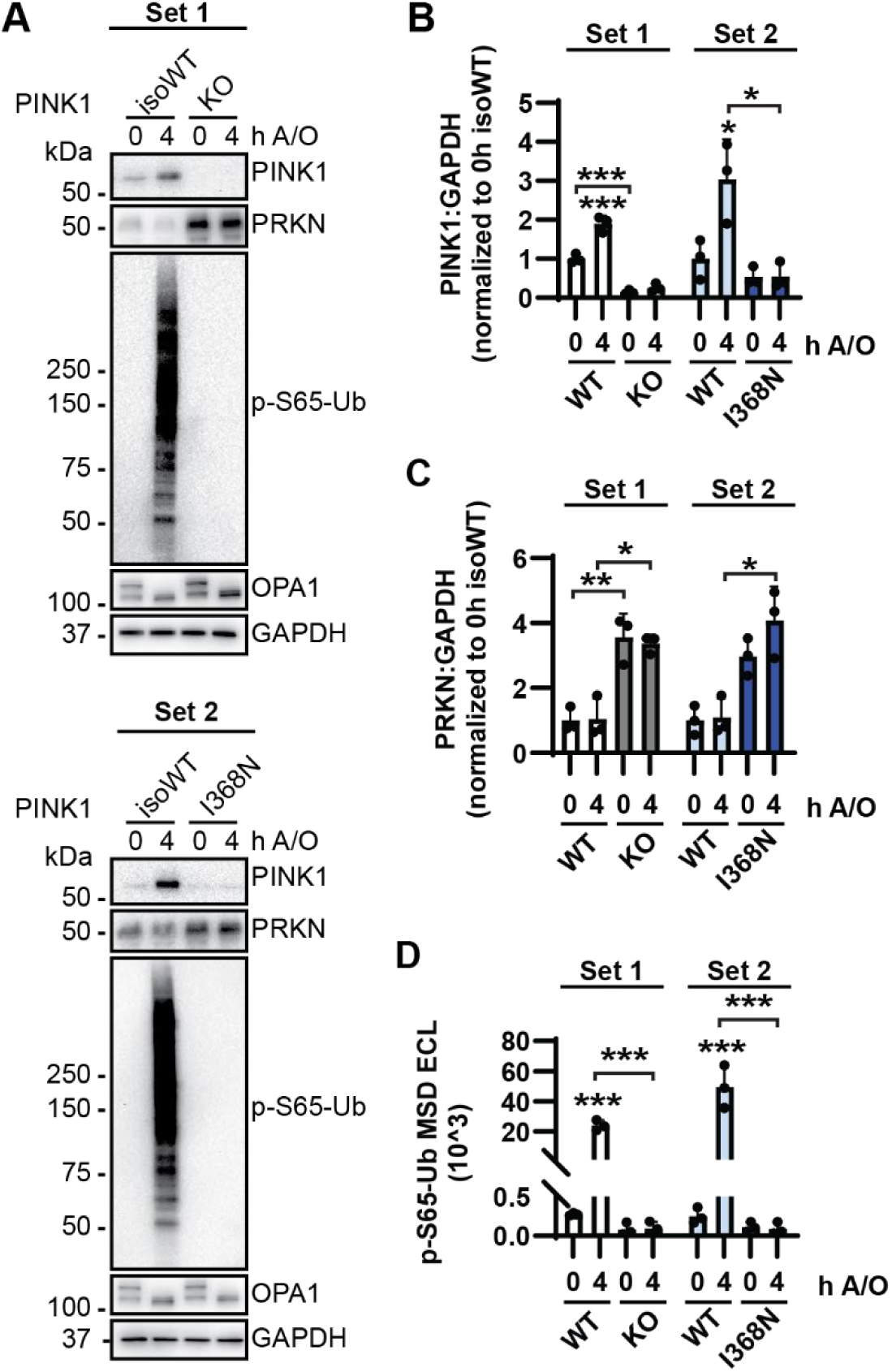
Loss of PINK1 activity leads to PRKN accumulation and the absence of p-S65-Ub signal in astrocytes. (**A**) Representative western blots for each set of iAS show levels of PINK1, PRKN, p-S65-Ub and OPA1 under baseline conditions and in response to A/O treatment. GAPDH was used as the loading control. Densitometric analysis of **(B)** PINK1 and **(C)** PRKN western blots shown under (a) and data points displayed as PINK1 or PRKN divided by GAPDH. **(D)** Quantification of p-S65-Ub levels measured by MSD sandwich ELISA. PINK1:GAPDH and PRKN:GAPDH data is represented as the PINK1 mutant iAS normalized to their corresponding 0h isoWT control set to 1, while MSD results are displayed as background corrected raw data. The mean of three independent differentiation experiments for each iAS set ± SD is depicted. Statistical analysis was performed using Two-Way ANOVA with Tukey’s multiple comparison test (***p<0.001, **p<0.01, *p<0.05). Asterisks on top of data points indicate individual comparisons to respective WT controls without PINK1 mutation.

### Loss of PINK1 reduces astrocyte reactivity and boosts pro-inflammatory signaling

Next, we assessed iAS purity to confirm whether loss of PINK1 activity hinders astrocyte differentiation. About 100% of astrocytes from both sets displayed positive signal for the stable astrocyte marker ALDH1L1, with no significant differences in signal intensity (**Fig 3, A to C**), confirming no influence of PINK1 on differentiation to astrocytes. Interestingly, both the PINK1 KO and PINK1 I368N iAS displayed significant reductions in the astrocyte reactivity markers GFAP and S100B relative to their respective isoWT controls (**Fig. 3, D to E**). To understand whether the reduction in intracellular GFAP and S100B signal could be the result of increased secretion, levels of both proteins were measured in iAS conditioned media from both sets of cells. Prior studies have shown that GFAP can be released extracellularly and detected in the bloodstream following acute CNS injury*^(29)^*. Yet, no significant differences in GFAP secretion were observed here (**Fig. 3F**). Although S100B can be secreted under healthy physiological conditions, excessive secretion is neurotoxic and a sign of injury, leading to dysregulation of neuronal firing via inhibition of potassium channels*^(30–32)^*. Interestingly, significant increases of S100B secretion were observed in PINK1 KO iAS, which showed a similar trend in PINK1 I368N iAS compared to their respective isoWT controls (**Fig. 3G**). However, increased secretion of these markers could not be explained by astrocyte death, as there were no differences in viability across genotypes (**Fig. 3, H to I**).

**Fig. 3.**
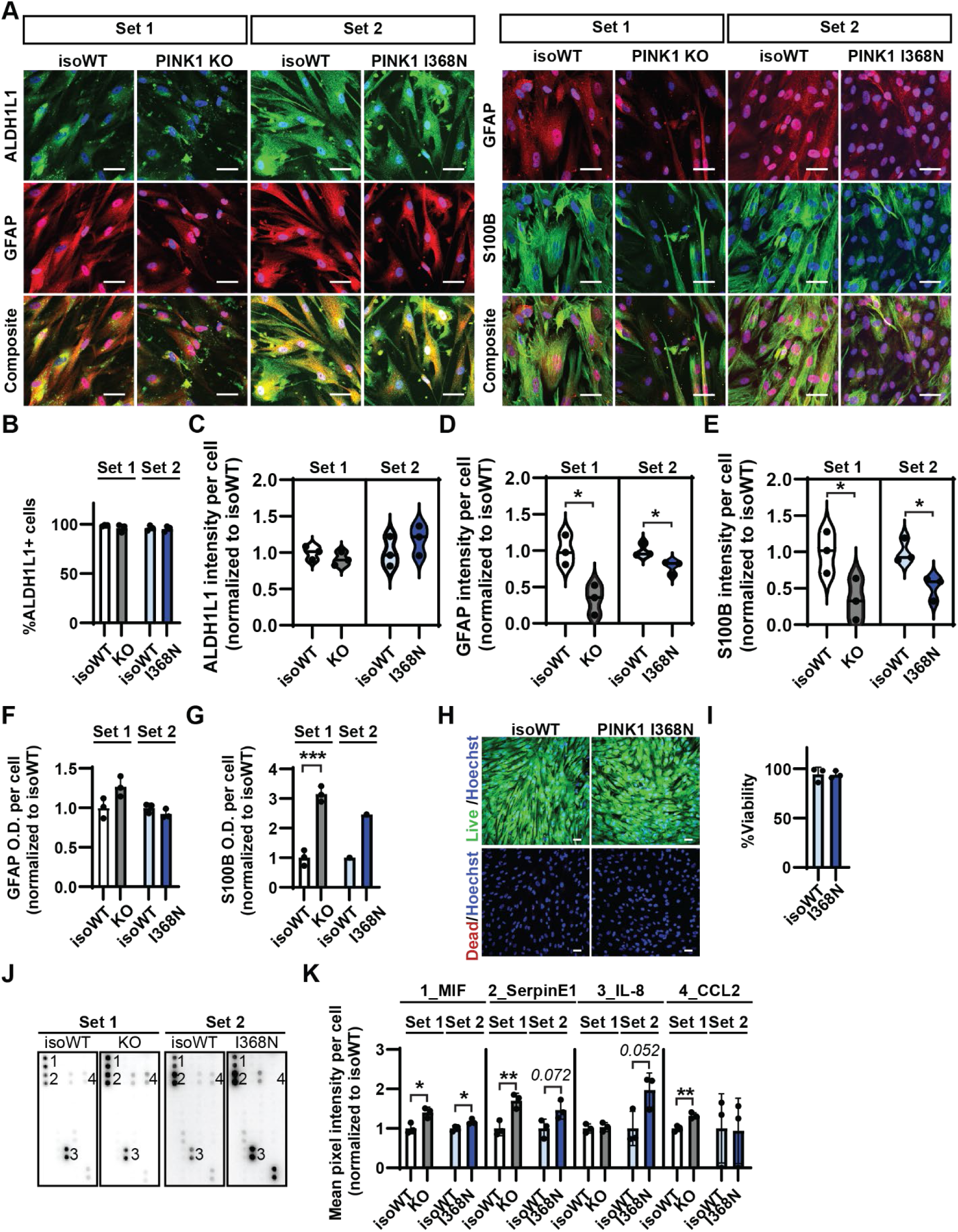
Loss of PINK1 reduces astrocyte reactivity and triggers release of pro-inflammatory danger signals. **(A)** Representative immunofluorescent images displaying about **(B)** 100% purity for the pan-astrocyte marker ALDH1L1 and **(C)** no significant differences in ALDH1L1 intensity across genotypes from both sets. Co-staining of GFAP and S100B revealed **(D-E)** significant decreases in signal intensity of both markers across both PINK1 mutants compared to their respective isoWT controls. Levels of secreted **(F)** GFAP and **(G)** S100B from astrocyte conditioned media were measured via sandwich ELISA. Average optical density (O.D.) normalized to cell count was quantified. **(H)** Representative immunofluorescent images using LIVE/DEAD assay and **(I)** quantification displaying no significant differences in cell viability (Live/Live+Dead)x100. Live cells are labeled by calcein-AM, while dead cells are labeled by ethidium homodimer-1. **(J)** Proteome profiler human cytokine array of iAS conditioned media. 1-4 labels represent significantly different markers in one or both iAS sets. **(K)** Quantification of dot blots is displayed as the mean pixel intensity of each marker normalized to a positive loading control of the respective isoWT. The data was then further normalized to the cell count for each respective genotype. Data is represented as the PINK1 mutant iAS normalized to their corresponding isoWT control set to 1. The mean of three independent differentiation experiments for each iAS set ± SD is depicted. PINK1 I368N and isoWT S100B sandwich ELISA represent N=1. Data was analyzed by using Student’s T-test (***p<0.001, **p<0.01, *p<0.05). Nuclei were stained with Hoechst (blue). Scale bar = 50 μm.

The secretomic profiles of the PINK1 mutant iAS were next unbiasedly defined using the Proteome Profiler Human Cytokine Array. Both the PINK1 KO and PINK1 I368N iAS secreted significantly more Macrophage Migration Inhibitory Factor/MIF compared to their respective isoWT (**Fig. 3, J to K**). The PINK1 KO iAS also displayed greater secretion of SerpinE1/PAI-1 and CCL2/MCP-1, while the I368N iAS showed a borderline significant increase in IL-8. Collectively, these markers are pro-inflammatory danger signals that are secreted by astrocytes to recruit microglia and immune cells to sites of injury and can promote neurotoxicity*^(33–37)^*. These findings indicate that loss of PINK1 activity in astrocytes increases pro-inflammatory signaling under baseline/unstimulated conditions, i.e., in the absence of dead or dying neurons that are known to trigger such response.

### Transcriptomic profiling of PINK1 I368N astrocytes

To uncover the mechanisms underlying how loss of PINK1 alters iAS inflammatory signaling, unbiased transcriptomic profiling was performed using bulk short read RNA sequencing of the PINK1 I368N iAS patient line and the corresponding isoWT control. Enrichment analysis confirmed that gene expression profiles of the cultured iAS resemble human astrocytes compared to other CNS cell types (i.e. compared to neurons, oligodendrocytes, endothelial cells, and monocytes) (**Fig. S2A**). After adjusting for batch effect and following normalization and dispersion estimation, differential gene expression analysis was performed using DESeq2, and clustering based on genotype via principal component analysis was confirmed (**Fig. S2, B to C**).

Differential gene expression analysis yielded 401 significantly different protein coding genes with 308 upregulated and 93 downregulated in the PINK1 I368N iAS compared to isoWT controls (adj. p value < 0.05, |Log2 FC| ≥ 1) (**Fig. 4A**). As expected, there were no changes in PINK1 expression in the PINK1 I368N kinase dead mutant compared to isoWT(*24*). Astrocyte-related genes ALDH1L1, GFAP, and S100B were also not among the differentially expressed genes (DEGs), further confirming that loss of PINK1 does not hinder astrocyte differentiation and indicating that reduction of intracellular GFAP and S100B protein levels are not due to loss of their gene expression.

**Fig. 4.**
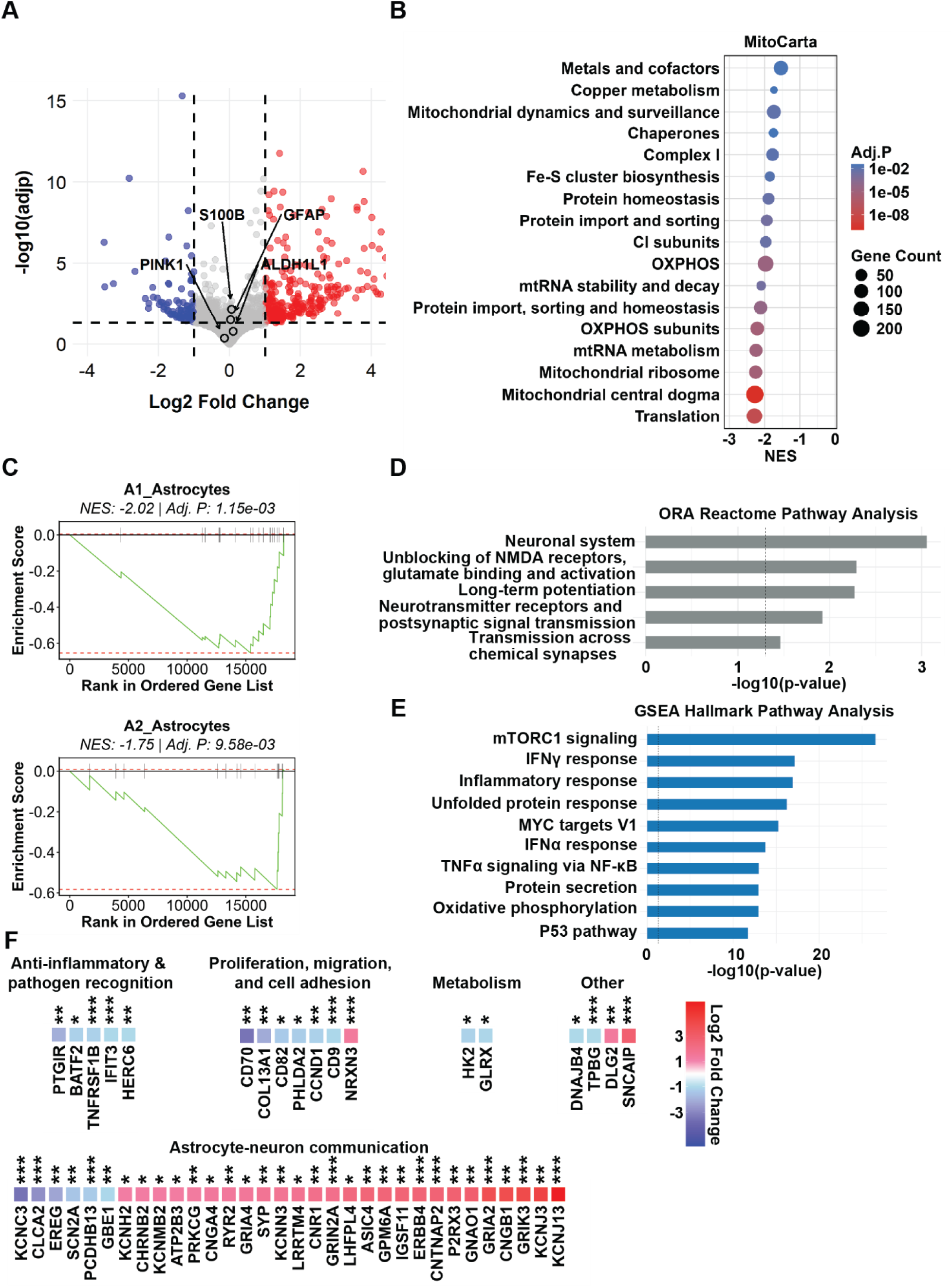
Transcriptional profiling of PINK1 I368N iAS. **(A)** Volcano plot displaying 401 differentially expressed genes (adj p < 0.05, |log2 FC| ≥ 1). Blue dots and red dots indicate downregulation and upregulation of genes in the PINK1 I368N compared to isoWT, respectively. Grey dots = not significantly different genes. Enrichment analysis was performed for comparison to **(B)** mitochondrial pathways via MitoCarta and **(C)** A1 and A2 transcriptomic signatures. NES = normalized enrichment score. Pathway analysis was subsequently performed via **(D)** over-representation analysis using all DEGs and **(E)** gene set enrichment analysis (GSEA) using the entire dataset. Leading edge DEGs from the top 5 ORA pathways and top 10 GSEA pathways were categorized as playing a role in **(F)** anti-inflammatory signaling & pathogen recognition; proliferation, migration, and cell adhesion; metabolism; other, mainly playing a role in cancer processes; or astrocyte-neuron communication. Fold change and NES data is represented as a ratio of expression in PINK1 I368N in relation to isoWT. Samples from five independent differentiations were analyzed. Adjusted p-values (adjp) were calculated using the Benjamini-Hochberg correction for multiple testing (***adj p<0.001, **adj p<0.01, *adj p<0.05).

Inability to selectively degrade damaged mitochondria resulting from loss of PINK1 kinase activity is known to cause metabolic dysfunction via buildup of reactive oxygen species, respiratory failure, and mtDNA leakage, which may thereby trigger an innate immune response*^(38–41)^*. However, these effects are yet to be investigated directly in glia. Therefore, enrichment analysis of mitochondrial pathways using MitoCarta was performed, confirming pervasive suppression of mitochondrial maintenance and bioenergetic machinery in the PINK1 I368N iAS compared to isoWT (**Fig. 4B**).

Mitochondrial dysfunction has been linked to glial and immune cell exhaustion and altered astrocyte reactivity(*12, 14*). To define the reactive profile of the PINK1 mutant astrocytes, gene set enrichment analysis was performed to compare the iAS transcriptome to canonical A1 and A2 signatures*^(42, 43)^*. A1 signatures are characterized by excessive pro-inflammatory signaling while A2 promotes neuroprotection via anti-inflammatory signaling and upregulation of neurotrophic factors*^(42)^*. The PINK1 I368N iAS displayed downregulation of canonical A1 and A2 signatures (**Fig. 4C**), further confirming the link between mitochondrial dysfunction and altered glial reactivity. A1 and A2 signatures have been previously defined from astrocytes in the presence of neuronal damage, reactive microglia, or treatment with cytokine cocktails mimicking microglial reactivity*^(42, 43)^*. While these iAS may adopt an A1 signature in the presence of reactive microglia, these findings indicate that the A1/A2 binary classification may be an oversimplification*^(32, 44)^*, underscoring the need to study astrocyte dysfunction also under unstimulated conditions to identify cell-type-specific, disease-modifying targets. Pathway analysis was additionally performed to further define the mechanisms underlying astrocyte dysfunction.

Over-representation analysis (ORA) using protein coding DEGs was then performed via the Reactome database. This revealed five dysregulated pathways related to altered astrocytes support for neuronal firing and neurotransmission (**Fig. 4D**). These results were further corroborated by Metascape enrichment analysis, showing clustering of gene ontology terms associated with astrocyte support for neurotransmission, maintenance of Ca^2+^ homeostasis, and others (**Fig. S3A**). Transcriptional Regulatory Relationships Unraveled by Sentence-based Text (TRRUST) database was then utilized to predict potential upstream transcriptional regulators mediating these gene expression changes (**Fig S3B**); TRRUST analysis revealed enrichment of networks associated with regulation of inflammatory signaling and astroglial activation (LEF1, SP4, HDAC1, ESR1, SP1, CTNNB1, SP3, JUN, HDAC2), many via NF-κB signaling*^(45–49)^*. Other potential regulators from TRRUST were implicated in glutamate signaling (REST)*^(50)^* and proliferation (NANOG)*^(51)^*.

Next, the entire data set was used to perform GSEA pathway analysis via the Hallmark database. The top ten altered pathways included mTOR signaling, various inflammatory signaling pathways, MYC targets V1 related to proliferation, metabolic processes, and others (**Fig. 4E**). The functions of Leading Edge DEGs driving the top five and top ten significantly dysregulated pathways observed in ORA and GSEA, respectively, were unbiasedly summarized using Rentrez (**Table S1**). Interestingly, there was a general downregulation of genes associated with anti-inflammatory signaling and pathogen recognition (**Fig. 4F**). This was accompanied by reduced expression of genes regulating proliferation, migration, cell adhesion, and metabolism. However, the majority of these leading edge DEGs were associated with astrocyte-neuron communication, particularly those playing a role in maintaining Ca^2+^ homeostasis and astrocyte mediation of excitatory neuronal signaling (**Fig. 4F**). Coupled with secretion of pro-inflammatory danger signals, reduction of reactivity markers, and metabolic defects via MitoCarta, these findings suggest that loss of PINK1 may induce astrocyte exhaustion and homeostatic collapse, resulting in the inability to adequately support neurotransmission.

### Functional validation of RNA sequencing results

To determine whether these transcriptional signatures would translate into functional deficits, biochemical and cell biological assays were performed to corroborate key pathways identified in the RNA sequencing results. Using Fluo-4, AM, significantly reduced levels of intracellular Ca^2+^ were revealed in the PINK1 I368N iAS compared to isoWT (**Fig. 5A**). These results validate findings from ORA and Metascape in addition to the downregulation of S100B protein levels, which is a critical player in the maintenance of Ca^2+^ homeostasis. The PINK1 I368N iAS also displayed significantly increased levels of p-S536-p65 (**Fig. 5B**), a core subunit of NF-κB and marker of signaling activation. p65 is phosphorylated in the cytoplasm and subsequently rapidly translocation to the nucleus to promote transcriptional activation*^(52)^*. In addition, downregulation of pathways associated with proliferation were validated with an EdU assay that showed about a 50% reduction in proliferation in PINK1 I368N compared to isoWT controls (**Fig. 5C**). Finally, mTORC1 signaling was the most significantly altered pathway via GSEA. This pathway is known to be critical for regulation of autophagy*^(53)^*. The PINK1 I368N iAS displayed increased levels of p-S2448-mTOR compared to isoWT controls, while total mTOR levels remained unchanged (**Fig. 5, D to F**). This was accompanied by a borderline significant reduction in autophagy as measured through LC3-II (**Fig. 5G**).

**Fig. 5.**
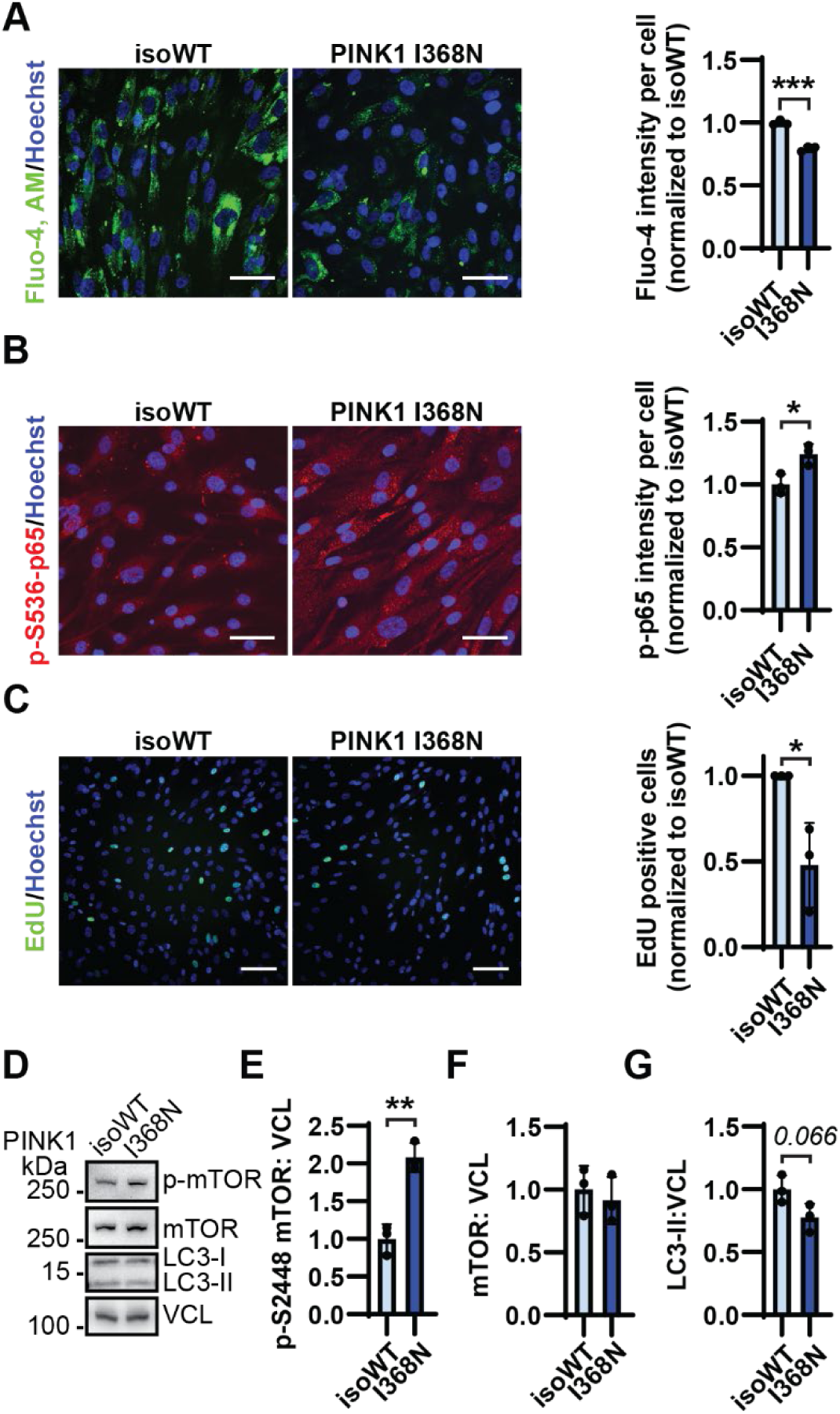
Functional validation of RNA sequencing results. Intracellular Ca^2+^ levels were labeled using **(A)** Fluo-4,AM and quantified as signal intensity normalized to cell count. **(B)** Representative immunofluorescent images and quantification of p-S536-p65 signal intensity normalized to cell count. Number of **(C)** EdU positive cells were quantified as normalized to cell count. **(D)** Representative western blot and quantification of **(E)** p-S2448-mTOR, **(F)** total mTOR, and **(G)** LC3-II levels in PINK1 I368N compared to isoWT controls divided by the loading control, vinculin (VCL). Data is represented as normalized to isoWT iAS set to 1. The mean of three independent experiments for each iAS set ± SD is depicted. Data was analyzed by using Student’s T-test (***p<0.001, **p<0.01, *p<0.05). Scale bar = 50 μm for Fluo-4, AM and p-S536-p65 experiments and 100 μm for EdU assay.

### mTOR inhibition rescues PINK1 mutant astrocyte pro-inflammatory signaling

Given both RNA sequencing results and biochemical validation at the protein level showing dysregulation to mTOR signaling, it was next investigated whether boosting autophagy via mTOR inhibition may compensate for loss of PINK1 mediated mitophagy. Rapamycin is an FDA approved mTOR inhibitor leading to autophagy activation and immunosuppression*^(54)^*. Therefore, we treated the PINK1 I368N iAS with rapamycin and confirmed successful mTOR inhibition via reduced p-S2448-mTOR without alteration of total mTOR levels, along with elevated LC3-II signal (**Fig. 6, A to D**). Rapamycin treatment was unable to rescue GFAP levels but did result in a borderline significant rescue of S100B (**Fig. 6, E to G**). This was accompanied by a rescue in secretion of MIF, which was observed previously upregulated in the PINK1 I368N compared to isoWT controls (**Fig. 6, H to I**). Importantly, the PINK1 KO iAS recapitulated these results. PINK1 KO iAS displayed upregulation of p-S2448-mTOR levels compared to isoWT that were successfully inhibited by rapamycin treatment (**Fig. S4A**). Following rapamycin treatment, the PINK1 KO iAS further displayed an increased trend in both GFAP and S100B levels (**Fig. S4, B to D**). This was also accompanied by a downregulation of secreted MIF and SerpinE1 (**Fig. S4, E to F**). Importantly, these data suggest that the astrocyte neuroinflammatory secretory phenotype resulting from loss of PINK1 may be pharmacologically reversible via boosting general autophagy.

**Fig. 6.**
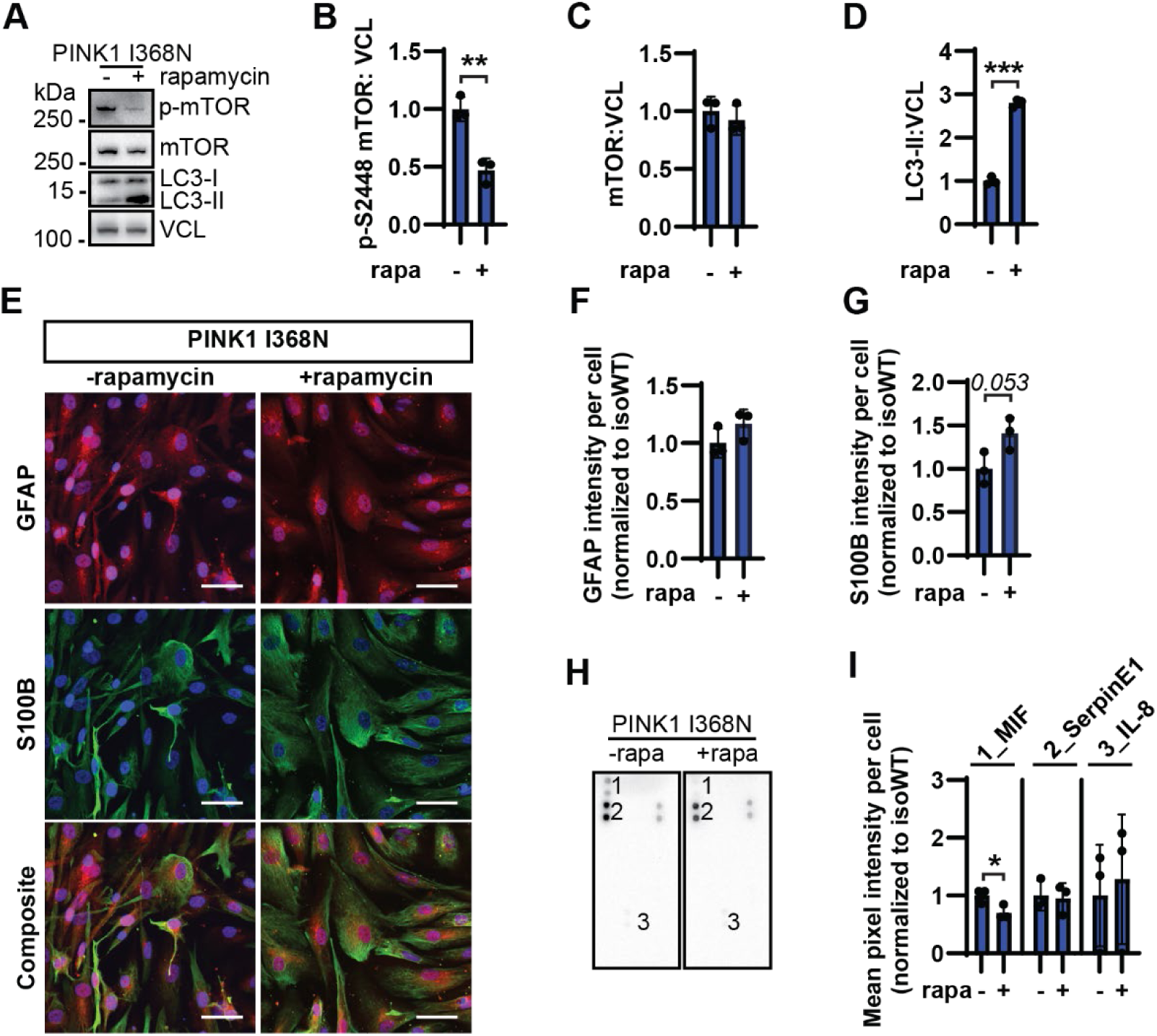
mTOR inhibition rescues pro-inflammatory signaling but not GFAP levels in PINK1 mutant iAS. PINK1 I368N iAS were treated with 500 nM rapamycin for one week. Treatment efficacy was confirmed via **(A)** representative western blot and quantification of **(B)** p-S2448-mTOR, **(C)** total mTOR, and **(D)** LC3-II levels divided by the loading control, VCL. **(E)** Representative immunofluorescence imaging and quantification of **(F)** GFAP and **(G)** S100B levels under baseline conditions and following rapamycin treatment. **(H)** Representative human cytokine array dot blot and **(I)** quantification displays rescue of MIF secretion in PINK1 I368N iAS in response to rapamycin treatment. Data is represented as normalized to isoWT iAS set to 1. The mean of three independent experiments for each iAS set ± SD is depicted. Data was analyzed by using Student’s T-test (***p<0.001, **p<0.01, *p<0.05). Scale bar = 50 μm.

### PINK1 I368N iAS reduce neuronal Ca^2+^ signaling and viability

Given our earlier findings that both sets of PINK1 mutant iAS increasingly secrete pro-inflammatory and potentially neurotoxic signals, we next aimed to elucidate the impact of the PINK1 mutant astrocyte secretome to neuronal health. To accomplish this, PINK1 I368N or isoWT iAS were co-cultured with PINK1 WT i3N for one week via transwell insert (**Fig. 7A**). The transwell system forms a physical barrier between the iAS and i3N, while permitting the flow of conditioned media through a porous membrane. Based on the findings that PINK1 mutant iAS secrete excessive amounts of S100B along with RNA sequencing results highlighting dysregulation of pathways associated with Ca^2+^ homeostasis, the i3N intracellular Ca^2+^ levels were measured. The i3N displayed significantly reduced levels of the intracellular Ca^2+^ indicator Fluo-4, AM, when cultured with the PINK1 I368N iAS compared to the isoWT iAS (**Fig. 7, B to C**). This was accompanied by ∼30% reduction in viability from the i3N cultured with the PINK1 I368N iAS (**Fig. 7, D to E**). A similar reduction in viabilty was seen when i3N were cultured with PINK1 KO iAS compared to isoWT iAS (**Fig. S5, A to B**). These findings indicate that PINK1 mutant astrocytes are direct contributors to neuronal damage and death, rather than acting solely as responders to neuronal injury in PD.

**Fig. 7.**
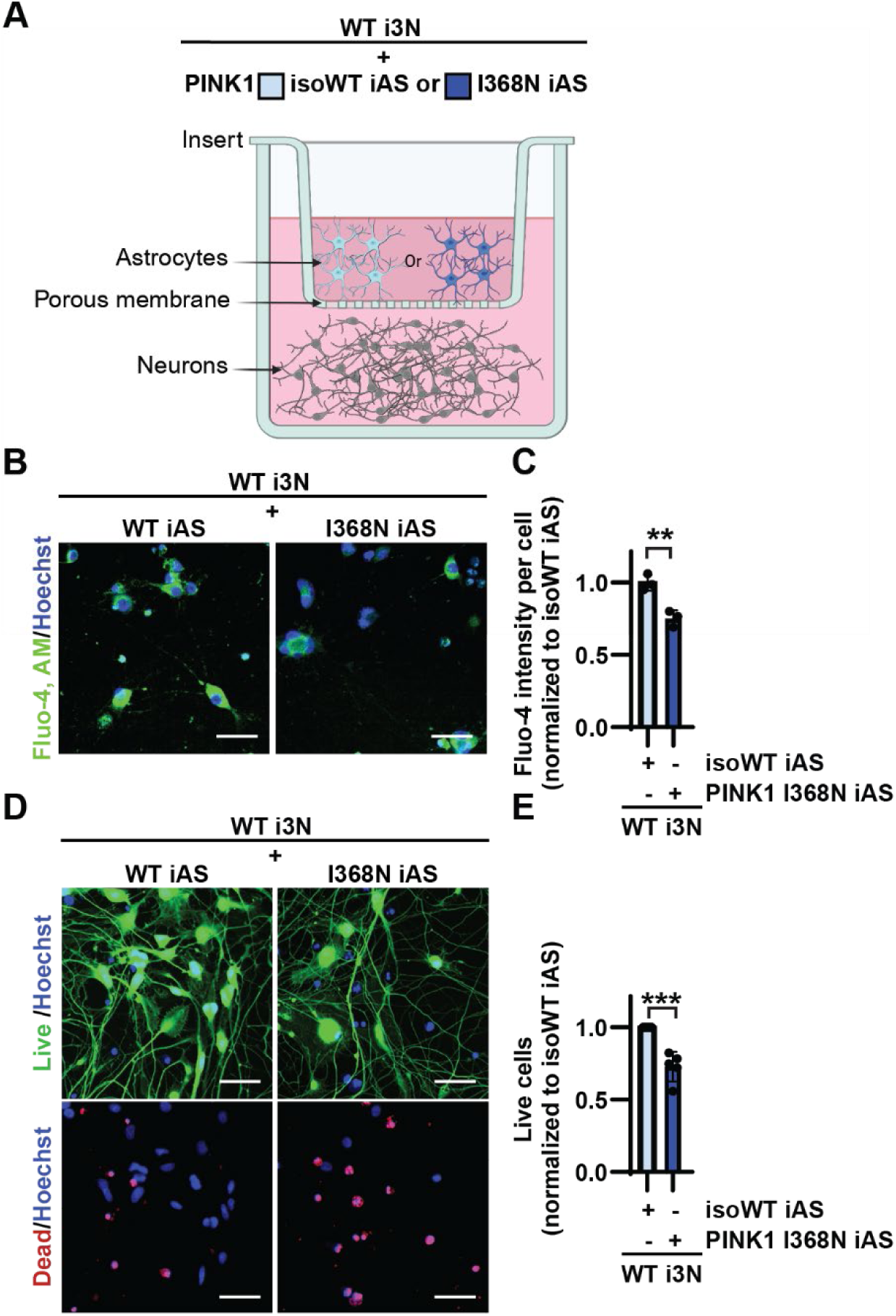
PINK1 mutant iAS secretome reduces neuronal Ca^2+^ levels and induces death. **(A)** Schematic illustration of experimental setup in which PINK1 I368N or isoWT iAS are seeded in a transwell insert, separated by a porous membrane from the WT i3N that are grown in the well, allowing the neurons to be exposed to the iAS secretome. Intracellular Ca^2+^ levels in the neurons were labeled using **(B)** Fluo-4, AM and **(C)** quantified as signal intensity normalized to cell count. Viability was measured through **(D)** immunofluorescent imaging using LIVE/DEAD assay and **(E)** quantified as the fraction of live cells. Data is represented as normalized to WT i3N cultured with isoWT iAS set to 1. The mean of three independent experiments for each iAS set ± SD is depicted in **(C)**, and as a ratio of mean viability in five independent experiments in **(E)**. Data was analyzed by using Student’s T-test (***p<0.001, **p<0.01). Scale bar = 50 μm. The transwell schematic was generated using BioRender.

## DISCUSSION

It is well known that dopamine neurons progressively degenerate in PD. While loss of PINK1 or PRKN function leads to their demise, our findings reveal that the ubiquitin kinase is also critical to the health and function of CNS cell types other than neurons. For the first time through this study, we defined the level of PINK1 and PRKN expression and activity in human astrocytes under baseline and acute stress conditions. We utilized two independent isogenic sets of gene-edited and patient-derived PINK1 mutant iAS to understand the importance of mitophagy signaling to astrocyte health and function at the transcriptomic, protein, and biological level. Collectively, these studies revealed that loss of PINK1 gene expression or enzymatic activity in astrocytes triggered release of neurotoxic and pro-inflammatory danger signals, reduced reactivity, impaired proliferative capacity, induced metabolic failure, and Ca^2+^ dyshomeostasis. Importantly, these events occurred in the absence of dead or dying neurons and without inflammatory activator treatment that could mimic the microenvironment stimulated by dying neurons or reactive microglia. Altogether, we revealed the importance of PINK1 signaling beyond neurons and that astrocytes should be considered potential drivers of disease onset rather than simple secondary reactors.

Although previous transcriptomic analysis from human brain has indicated that PINK1 and PRKN are ubiquitously expressed*^(5, 19)^*, their level of activity in non-neuronal cell types has remained understudied. Prior reports utilizing primary rodent astrocytes indicated increased levels of the PINK1 and PRKN product p-S65-Ub in response to mitochondrial stress in comparison to other cell populations, including cortical neurons*^(26, 27)^*. Here we demonstrate that human iAS and cortical-like i3N have indeed comparable levels of PINK1, PRKN, and p-S65-Ub under both baseline and acute mitochondrial stress conditions. These data indicate that PINK1-PRKN signaling is active in astrocytes, highlighting the need to better define the level of PINK1-PRKN signaling and their protective effects across cell types under physiological and relevant pathological conditions to determine respective susceptibilities.

Loss of PINK1 activity also reduced the levels of reactivity markers GFAP and S100B in cultured astrocytes, similar to observations from PRKN mutation carrier post-mortem human brain*^(20, 21)^*. Of note, this was accompanied by increased secretion of pro-inflammatory danger signals and NF-κB activation. S100B deficiency could result from its excessive secretion, which is a known activator of NF-κB signaling*^(30, 31)^*. Interestingly, PINK1’s enzymatic activity is a contributor to anti-apoptotic NF-κB activation, while its loss of function leads to maladaptive NF-κB responses*^(15, 55)^*. Prior studies already suggested that mitophagy is critical for immune signaling regulation and suppression*^(56–60)^*. Building on these insights, we observed that genetic inactivation of PINK1 significantly altered the astrocyte inflammatory profile and triggered NF-κB activation even in the absence of exogenous inflammatory stress. However, it remains to be known whether complete loss of PINK1 activity is required or if even reduced PINK1-PRKN signaling, as seen in sporadic PD, can trigger such inflammatory response*^(61, 62)^*. The therapeutic potential of boosting PINK1-PRKN mitophagy has led to multiple mitophagy activating drugs moving into clinical trials*^(63)^*. It will be critical to determine whether these mitophagy activators may rescue glial dysfunction induced by PINK1-PRKN deficits.

Our transcriptomic, biochemical, and functional studies further revealed defects to PINK1 mutant astrocyte proliferative capacity, metabolism, and Ca^2+^ homeostasis. Proliferation is a critical aspect of astrocyte reactivity, with its reduction implicated in slowed wound healing and exacerbated damage*^(64)^*. A previous study also observed proliferation defects in PINK1 deficient primary mouse astrocytes*^(65)^*. Metabolic deficits are a hallmark of PINK1 loss of function, characterized by mitochondrial dysfunction, reactive oxygen species production, and disruption to ATP synthesis*^(3, 66)^*, which was reflected in our enrichment analysis results. Prior studies have also identified that loss of PINK1-PRKN signaling leads to Ca^2+^ dyshomeostasis in Drosophila, iPSC-derived NPCs, and mouse embryonic fibroblasts, potentially due to increased ER calcium release or impaired mitochondrial Ca^2+^ uptake*^(67, 68)^*. Ca^2+^ signaling is vital to astrocytes as it regulates gliotransmission and is critical for support of neurotransmission and synaptic plasticity*^(69)^*. Consistent with the functional consequences of Ca^2+^ dyshomeostasis, our transcriptomic analysis highlighted marked alterations to pathways associated with regulation and support for neurotransmission in the PINK1 mutant iAS. Collectively, excessive secretion of pro-inflammatory danger signals coupled with reduced reactivity, impaired proliferative capacity, metabolic failure, and Ca^2+^ dyshomeostasis point to astrocyte homeostatic collapse and exhaustion. Indeed, glial and immune cell exhaustion have been previously linked to altered PINK1 levels and mitochondrial dysfunction*^(12, 70)^*.

GSEA revealed mTOR signaling as the most dysregulated pathway in the PINK1 I368N iAS compared to isoWT, which we subsequently validated at the protein level for both PINK1 I368N and KO iAS. The mTOR signaling pathway is critical for regulation of autophagy, while its pharmacological inhibition boosts autophagic activation and immunosuppression*^(54)^*. We therefore aimed to understand whether boosting autophagy activation via rapamycin, an FDA approved mTOR inhibitor, could compensate for loss for PINK1-PRKN signaling in human astrocytes. Indeed, rapamycin treatment rescued secretion of the pro-inflammatory danger signals. However, rapamycin is also known to reduce intracellular Ca^2+^ signaling and proliferation*^(71, 72)^*. Therefore, it remains to be known whether it can fully rescue astrocyte dysfunction induced by loss of PINK1 activity. Nevertheless, results of this study indicate that pro-inflammatory danger signals secreted by PINK1 mutant iAS are pharmacologically reversible, thereby opening doors for future investigation into therapeutic targeting of astrocytes in PD to mitigate neuronal damage.

Finally, our study unveiled that the PINK1 mutant iAS secretome alone was sufficient to impair neuronal Ca^2+^ levels and reduce neuronal viability by almost 30%. Following inflammatory stimulation, PINK1 KO primary mouse astrocytes have been shown to induce neuronal apoptosis in the presence of reactive microglia, or under conditions designed to simulate the reactive microglial environment*^(27, 73)^*. However, neuronal damage was not observed under baseline conditions. These contrasts between PINK1 mutant human and mouse cultures may be reflective of the fact that loss of PINK1-PRKN signaling in mice does not lead to neurodegeneration and PD-like symptoms without the addition of exogenous stressors or extreme aging*^(74)^* . A strength of our approach was the use of mixed genotypes, combining PINK1 mutant astrocytes with wild-type neurons, which enabled us to isolate astrocyte-specific effects and demonstrate non-cell-autonomous roles of PINK1 signaling.

While dopamine neurons are known to degenerate upon PINK1-PRKN pathway inactivation, the results of our study revealed that astrocytes are also vulnerable and directly induce neuronal death. While we did perform unbiased profiling of the PINK1 mutant iAS secretome, we do not propose that our list of inflammatory danger signals is all-encompassing or that it will remain unchanged in the presence of reactive microglia. In fact, we hypothesize that the observed astrocyte dysfunction would be exacerbated in human brain and lead to a vicious cycle, wherein neuroinflammation contributes to neuronal loss that subsequently amplifies inflammatory signaling. Furthermore, while our data show that the PINK1 mutant iAS secrete pro-inflammatory danger signals, it is also possible that they neglect to secrete neurotrophic/neuroprotective factors. These data lay the groundwork for future studies to utilize more complex models, such as human midbrain organoids, or sensitive methods, such as proteomic profiling of conditioned media or single cell RNA sequencing, to better understand the inflammatory and reactivity profiles triggered by loss of PINK1 in glia. Additionally, while this study provides critical mechanistic insight that loss of PINK1 enzymatic activity induces homeostatic collapse, exhaustion, and neurotoxic overload to human iAS, further investigation is needed to determine the exact order of events and to tease out the collective versus independent contributions of PINK1 and PRKN in astrocyte health and function. Finally, the results of this study have vital implications for guiding development of disease modifying therapies, as targeting both nigrostriatal dopaminergic neurons and glia, rather than neurons alone, may be necessary to halt disease progression. Overall, our findings unveil the critical and urgent need to further study the impact of PINK1 loss of function on cell types beyond neurons, and the role of glial-neuron crosstalk in PD pathogenesis.

## MATERIALS AND METHODS

### iPSC culture and differentiation

PINK1 KO (ASE-9402) iPSCs were generated via CRISPR-Cas9 gene editing to induce compound heterozygous frameshift variants in PINK1 exon 2 from a Caucasian male parental isogenic control (isoWT) iPSCs (ASE-9109) and purchased from Applied Stem Cell. PINK1^I368N^ iPSCs, derived from a Caucasian female EOPD patient, and corresponding isogenic gene-corrected control (isoWT) iPSCs were obtained from the NINDS Human Cell and Data Repository (NHCDR, ND50100, ND50012). WT KOLF2.1J iPSCs were purchased from Jackson Laboratory (JIPSC001492). All iPSCs were tested for normal karyotype and pluripotency, and negative for mycoplasma contamination. Undifferentiated iPSCs were maintained on a Matrigel matrix (Corning, 354277) coated surface diluted in DMEM/F12 (Gibco, 10565018) and fed with mTeSR Plus (Stemcell technologies, 100–0276). Full media changes were performed every other day or when the media color turned yellow.

### NPC differentiation

iPSCs were first differentiated into NPCs and then astrocytes, following an established protocol*^(23)^*. In brief, once the iPSCs reached 75% confluency, they were dissociated using Accutase (Millipore, SCR005) and plated in STEMdiff SMADi Neural Induction media (StemCell, 08581) with 10 µM Y-27632 ROCK inhibitor (Selleckchem, S1049) at 1.92 x 10^6^ cells/well into a Matrigel-coated 6-well plate. Full media changes were performed every day. Every, 6-9 days, the cells were dissociated using Accutase and plated at the same density in media with 10 µM ROCK inhibitor. Around 14-18DIV of differentiation, the cells were again dissociated and plated into a 96-well plate at 5 x 10^4^ cells/well for immunofluorescent imaging to confirm NPC purity.

### iAS differentiation

After confirming NPC purity, the cells were seeded into a 6-well plate at 1.6 x 10^5^ cells/well for astrocyte differentiation using STEMdiff SMADi Neural Induction Media without ROCK inhibitor. Twenty-four hours later, a full media change was performed using Astrocyte Medium (ScienCell, 1801), containing 2% fetal bovine serum (FBS) (ScienCell, 0010). Full media changes were performed three times per week. The cells were dissociated using Accutase every 9 days or upon reaching 90-100% confluency and plated into 6-well plates at 1.6 x 10^5^ cells/well. On day 30 of differentiation, the cells were dissociated using Accutase and plated into 96-well plates at 5 x 10^4^ cells/well and 6-well plates at 1.6 x 10^5^ cells/well using Astrocyte Medium without FBS. Full media changes were performed every other day. All experiments were performed in the absence of FBS for at least 7 days, with the exception of EdU, which was performed upon withdrawing FBS for 24 hours.

### Differentiation of iPSCs to i3N

WT iPSCs with inducible expression of neurogenin-2 by doxycycline were purchased from Alstem (iP11N). The cells were differentiated as previously described*^(25)^*. In brief, once the iPSCs reached 75% confluency, they were dissociated using Accutase and seeded at 1 × 10^6^ cells/well onto a Matrigel-coated 6-well plate using KnockOut Dulbecco’s Modified Eagle Medium (DMEM)/F12 (Thermo Fisher Scientific, 12660012) containing 1% *N*-2 supplement (Thermo Fisher Scientific, 17502048), 1% Glutamax (Thermo Fisher Scientific, 35050061), 1% MEM Non-Essential Amino Acids (NEAA) solution (Thermo Fisher Scientific, 11140050), 2 µg/mL doxycycline (Applichem, A2951), and 10 µM Y-27632 ROCK inhibitor. Full media changes were subsequently performed for the next 3 days without ROCK inhibitor, and doxycycline was freshly supplemented to the media each day. On day 4, the cells were again dissociated using Accutase and transferred onto a 0.1 mg/ml poly-L-ornithine/PLO (Sigma Aldrich, P3655)-coated 96-well plate at 2.3 x 10^4^ cells/well using KnockOut DMEM/F12 and Neurobasal A Medium (Thermo Fisher Scientific, 10888022) containing 0.5% *N*2 supplement, 1% B27 supplement (Thermo Fisher Scientific, 17504044), 1% Glutamax, 1% NEAA, 1 µg/ml laminin, 10 ng/mL BDNF (brain derived neurotrophic factor; Peprotech, 450–02), 10 ng/ml NTF3 (neurotrophin 3; Peprotech, 450–03), and 2 µg/ml doxycycline. At D14 of differentiation, PINK1 I368N or isoWT iAS were seeded into a Matrigel matrix-coated 96-well transwell system (Corning, 3380) at 1.15 x 10^4^ cells/well and experiments were performed 1 week following iAS seeding. All cells were maintained at 37°C and 5% CO_2_ in a humidified atmosphere.

### Cell culture treatment

To induce mitochondrial depolarization, the cells were treated for 4 hours with a mixture of 4 µM antimycin A (Sigma, A8674) and 10 µM oligomycin A (Sigma, 75351). For rapamycin rescue experiments, full media changes were performed three times within seven days with 500 nM rapamycin (Selleckchem, S1039).

### Cell lysis and western blot

Cells were washed two times with cold PBS (Boston Bioproducts, BM-220) and harvested in RIPA buffer (50 mM Tris [Sigma-Aldrich, 648311], pH 8.0, 150 mM NaCl [Sigma Aldrich, S5886], 0.1% SDS [Fisher Scientific, BP166-500], and 0.5% Deoxycholate [Sigma-Aldrich, D6750], 1% NP-40 [Sigma-Aldrich, I3021]), supplemented with a protease (Sigma-Aldrich, 11697498001) and phosphatase inhibitor (Sigma-Aldrich, 04906837001) cocktail. The samples were centrifuged at 21,000× *g* for 15 minutes at 4 °C, supernatant was collected, and protein concentration was measured by bicinchoninic acid (Thermo Fisher Scientific, 23225). 10µg of the lysates were loaded onto 8–16% Tris-Glycine gels (Thermo Fisher Scientific, XP08165BOX) or 16% Tris Glycine gels (Thermo Fisher Scientific, XP00165BOX) and transferred onto polyvinylidene fluoride membrane (PVDF) membranes (Millipore, IEVH00005). Membranes were then blocked with 5% skim milk in TBS with 0.1% Tween (TBST) for 1 h at RT and incubated with primary antibodies overnight at 4°C.

**Table 1:**
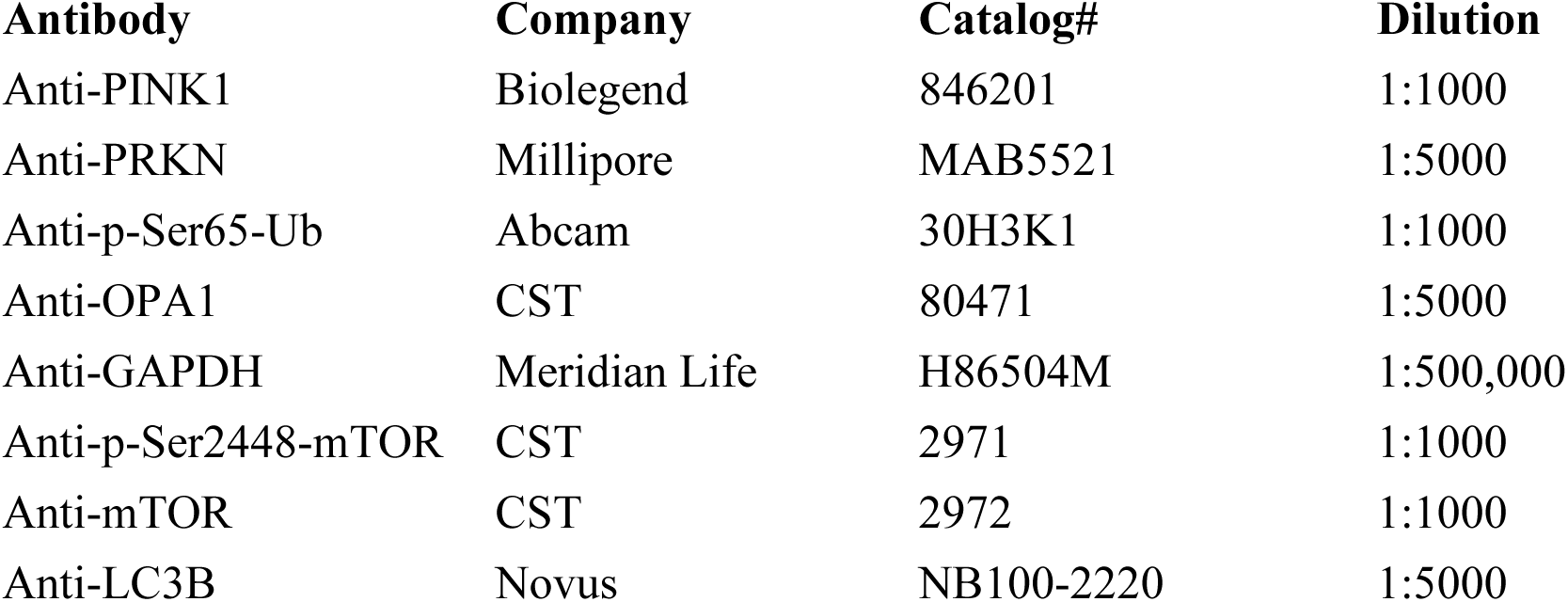
Antibodies used for immunoblotting.

Membranes were incubated for 1 h at room temperature in secondary antibodies. The following secondary antibodies from Jackson ImmunoResearch (West Grove, PA, USA) were used: donkey anti-rabbit IgG (11-035-152), goat anti-mouse IgG1 (115-035-205), and goat anti-mouse IgG2b (115-035-207). Signal was visualized using Immobilon Western Chemiluminescent HRP Substrate (Millipore Sigma, WBKLS0500) and recorded with a Chemidoc MP imaging system (Bio-Rad, Hercules, CA, USA).

### MSD ELISA

MSD ELISA against p-S65-Ub was performed as previously described*^(28)^*. Briefly, 96-well plates (Meso Scale Diagnostics, L15XA–3) were coated with capture antibody in 200 mM sodium carbonate buffer pH 9.7 overnight at 4°C. The next day, the wells were washed 3 times with TBST wash buffer and subsequently blocked for 1 h with blocking buffer (1% BSA [Boston BioProducts, Inc., *P*-753] in TBST). 5 µg of cell lysates diluted in blocking buffer and RIPA were loaded to the plates in duplicates at RT for 2 h on a shaking platform at 600 rpm. The wells were washed 3 times with TBST wash buffer and then detecting antibody was added for 2 h at room temperature. The wells were again washed 3 times and incubated with sulfo-tag labeled secondary antibody (Meso Scale Diagnostics, R32AC–1) for 1 h. Following a final 3 washes, the signal was measured following the addition of MSD Gold Read buffer (Meso Scale Diagnostics, R92TG–2) on a MESO QuickPlex SQ120 (Meso Scale Diagnostics, Rockville, MD, USA).

### Immunofluorescence

The cells were fixed using 4% paraformaldehyde (Sigma, 441244) for 15 minutes followed by 3 washes with PBS. The cells were permeabilized with 0.1% Triton X-100 (Sigma, X100) in PBS for 15 minutes and subsequently blocked using a mixture of 5% goat serum (Invitrogen, 16210072) + 5% BSA in PBS for 1 hour. The cells were incubated with primary antibodies diluted in blocking buffer overnight at 4°C.

**Table 2:** Antibodies used for immunofluorescence imaging.

| Antibody | Company | Catalog# | Dilution |
| --- | --- | --- | --- |
| Anti-Oct-4 | Novus | NB100-2379 | 1:100 |
| Anti-Nanog | Abcam | ab173368 | 1:100 |
| Anti-Nestin | Novus | NB100-1604 | 1:100 |
| Anti-Pax6 | Biolegend | 901302 | 1:100 |
| Anti-ALDH1L1 | Thermo | 14-9595-82 | 1:100 |
| Anti-GFAP | Aves Lab | GFAP857997 | 1:100 |
| Anti-S100B | Sigma | S2532 | 1:100 |
| Anti-TUBB3 | Novus | NB100-1612 | 1:500 |
| Anti-p-Ser536-p65 | CST | 3033 | 1:500 |

The next day, the cells were rinsed 3 times with PBS and then incubated with secondary antibodies for 1 h at RT. The following secondary antibodies from Molecular Probes was used: goat anti-rabbit 568 (A11011), goat anti-mouse 488 (A11001), goat anti-chicken 568 (A11041), goat anti-rabbit 488 (A11034), goat anti-chicken 647 (A21449). The cells were rinsed with PBS and then incubated for 10 minutes with Hoechst (Invitrogen, H21492) followed by a final PBS rinse.

Plates were imaged using an Operetta CLS High Content imaging system (Revvity, Waltham, MA, USA) with a 20× water immersion objective using a 2 × 2montage (no gaps) per well. Purity analysis was performed using ImageJ analysis software Cell Counter plugin. In each experiment, three fields of view from three replicate wells were quantified, with approximatively 1000 cells analyzed per genotype. Intensity analysis was performed using Harmony software. In each experiment, nine fields of view from three replicate wells were quantified, with approximately 3000 cells analyzed per genotype. Three independent experiments were performed for all assays.

### S100B and GFAP ELISAs, and protein array

Following a 3-day incubation of the cells with Astrocyte Medium, particulates were removed by centrifugation for 5 minutes at 500g at 4°C. The supernatant was then stored in aliquots and flash frozen to prevent multiple freeze thaw cycles. The levels of secreted GFAP were tested using the Human GFAP DuoSet ELISA Kit according to manufacturer’s guidelines (R&D Systems, DY259405). The levels of secreted S100B were tested using the Human S100 Beta ELISA kit according to manufacturer’s guidelines (Proteintech, KE00103). Data was analyzed by calculating the average optical density per sample loaded in duplicate and subtracting the average of a zero-standard absorbance.

The levels of cytokines and chemokines were screened in the cell culture supernatant using the Proteome Profiler Human Cytokine Array (R&D Systems, ARY005B), according to manufacturer’s guidelines. Equal volume of conditioned media was loaded onto each membrane per experiment. The signal of the protein array’s dot blots for each PINK1 mutant and corresponding isoWT control were recorded side-by-side to control exposure time using a Chemidoc MP imaging system (Bio-Rad, Hercules, CA, USA). The signal intensity was quantified using Image Studio software. The average intensity of duplicates was calculated and data across experiments were normalized according to the average of 4 reference signal controls that are included on each dot blot. To account for differences in proliferation between PINK1 mutant and isoWT iAS, the data was further normalized to the number of cells.

### LIVE/DEAD assay

The percentage of live cells was determined during the LIVE/DEAD Viability/Cytotoxicity Assay kit (Thermo Fisher Scientific, L32250) following the manufacturer’s protocol. The cells were incubated with the LIVE/DEAD and Hoechst reagents for 30 minutes at RT. The cells were then immediately imaged using an Operetta CLS High Content imaging system with a 20× water immersion objective using a 2 × 2montage (no gaps) per well. The percentage of live divided by total cells were quantified using the ImageJ Cell Counter.

### RNA sequencing and pathway analysis

Five independent batches of differentiated PINK1 I368N and isoWT control iAS were cultured. RNA was isolated from a total of 5 PINK1 I368N and 5 isoWT control iAS samples using the RNeasy Mini Kit (Qiagen, 74104). Samples were sequenced using the IlluminaNovaSeqX+10B using paired end mode at the Mayo Clinic Genome Analysis Core in Rochester, MN. Sequencing reads were aligned to the reference genome hg38, and raw gene read counts were generated using the Mayo Clinic RNA-seq analytic pipeline, MAP-RSeq(*75*). On average, 99% of reads from each sample mapped to the reference genome.

To analyze enrichment between iAS and human cortical CNS cell types from GSE73721 (Zhang et al., 2016 dataset), we followed methods outlined by similar studies*^(23)^*. In brief, both datasets were combined, adjusted for library size by counts per million (CPM), and a log transformation was applied. Effect size was determined using Spearman’s rank correlation coefficient, and the enrichment heatmap was plotted based on Z-score scaling.

For DEG analysis, principal components analysis was performed and did not reveal any outlier samples to be excluded in downstream processing. Source of variation analysis was conducted using the variancePartition R package (v.1.38.2) to assess the contribution of technical covariates explaining gene expression variation to determine covariate selection for the DEG model. RNA integrity number (RIN) was not a substantial contributor to source of variation, as RIN values ranged from 9.9-10. False Discovery Rate was used to adjust p values. Differential expression analysis was conducted using the R package DESeq2(*76*) v1.49.4, including batch as a covariate. We stabilized log2-fold change estimates across the dynamic range of expression via approximate posterior estimation for generalized linear model (apeglm) shrinkage(*77*). DEGs were defined by |log2 FC| ≥ 1 and adjusted p-value < 0.05. We observed 573 DEGs across all biotypes. We further filtered by protein coding genes, leading to 401 DEGs.

Gene set enrichment analysis (GSEA) was performed using the fgsea R package v1.34.2 for Hallmark(*78*) enriched Gene Ontology (GO), A1 and A2 astrocyte signatures, and MitoCarta v3.0(*79*). A1 (HLA-E, C3, SERPING1, HLA-B, HLA-C, HLA-F, LIG1, GBP2, FBLN5, SRGN, AMIGO2, PSMB8, HLA-A, ISG15, IFITM3, OAS1, STAT1, CXCL10, ICAM1) and A2 (CLCF1, TGM1, PTX3, S100A10, SPHK1, CD109, PTGS2, EMP1, SLC10A6, TM4SF1, B3GNT5, CD14, CLU, CD14, SPARCL1, SPP1, LIF) signatures were classified using well-defined and established datasets*^(42, 43)^*. Gene ranks were performed using the Wald statistic.

ORA was performed for all protein coding DEGs using the Reactome annotation gene set using ReactomePA R package v1.52.0. The biomaRt R package v2.64.0 was used to map Ensembl gene identifiers to gene symbols. Leading edge DEGs from GSEA and ORA were extracted and references were unbiasedly identified using the rentrez function via R script. Rentrez works to search the NCBI database to find records that match key words. Key word “astrocyte” and gene of interest were used to find records and subsequently selected based on relevance and most recent publication date. For cases in which no references were found in relation to astrocytes, the function of the gene itself was described.

Metascape enrichment analysis was performed using all protein coding DEGs. The network of enriched terms was plotted based on cluster ID. TRRUST was performed via Metascape. All plots, with the exception of those generated through Metascape, were produced using ggplot2 v4.0.2.

### Ca^2+^ signaling assay

Astrocytes were incubated with 2 µM Fluo-4, AM (Thermo Fisher Scientific, F14201) for 30 minutes at 37°C and 5% CO_2_ in a humidified atmosphere. The cells were then washed three times with HEPES solution (Sigma, H3537). On the second wash, the cells were incubated for 10 minutes with Hoechst (Invitrogen, H21492). The cells were then allowed to recover for 25 minutes at RT before imaging using the Operetta CLS High Content imaging system with a 20× water immersion objective using a 2 × 2montage (no gaps) per well. Harmony analysis software was used to make a cell mask to quantify the average intensity of Fluo-4, AM signal normalized to total number of cells per well.

### EdU Assay

As previously described, the astrocytes were re-plated at day 30 of differentiation using Astrocyte Medium without FBS. Within 24 hours after re-plating, proliferation was measured using the EdU Staining Proliferation Kit (Abcam, ab222421), according to the manufacturer’s guidelines. Plates were imaged using the Operetta CLS High Content imaging system with a 20× water immersion objective using a 2 × 2montage (no gaps) per well. The number of EdU positive cells were quantified using the ImageJ Cell Counter plugin.

### Statistics

All statistical analyses were performed with GraphPad Prism software or R. Details including sample sizes and statistical tests are described in figure legends. Results are displayed as mean ± SD. Results were compared using either a two-sided Student’s t-test or a two-way ANOVA followed by two-sided Tukey’s multiple comparisons test. P < 0.05 was considered statistically significant. For the RNA sequencing results, raw p-values were calculated using the two-sided Wald test followed by Benjamini-Hochberg FDR for multiple testing correction. Adjusted p value < 0.05 and |log2 fold change| ≥ 1 was used as the significance cutoff.

## Data availability

All data generated or analyzed during this study are included in this published article and supplementary data files. Other requests for data used in this manuscript can be addressed to the corresponding author and are available upon request.

## Acknowledgments

We are grateful to all patients and patients’ family members for donation and participation in research studies. We would like to thank our collaborator Dr. Takahisa Kanekiyo for his expertise and guidance in selecting protocols for differentiation of the iPSCs into NPCs. Bulk RNA sequencing and subsequent primary and secondary computational analyses were performed by the Mayo Clinic Genomics Core. We thank all members of the Springer lab for helpful discussions.

## Author contributions

**G.F.:** Manuscript writing, conceptualization, designed and analyzed experiments, data interpretation. **D.N.D.:** Advised and assisted with experimental design, data collection, and data interpretation. **D.H.H.**: Assisted with data collection. **O.A.R**.: Advised on experimental design and data interpretation. **F.C.F.**: Manuscript editing, guidance with experimental design, analysis, and data interpretation. **W.S.**: Manuscript writing and editing, guidance with experimental design, analysis, and data interpretation. Funding acquisition and supervision.

