## Supplementary Figures and Tables for "PINK1 loss in astrocytes triggers inflammatory dysfunction and neuronal death"

Fig. S1.

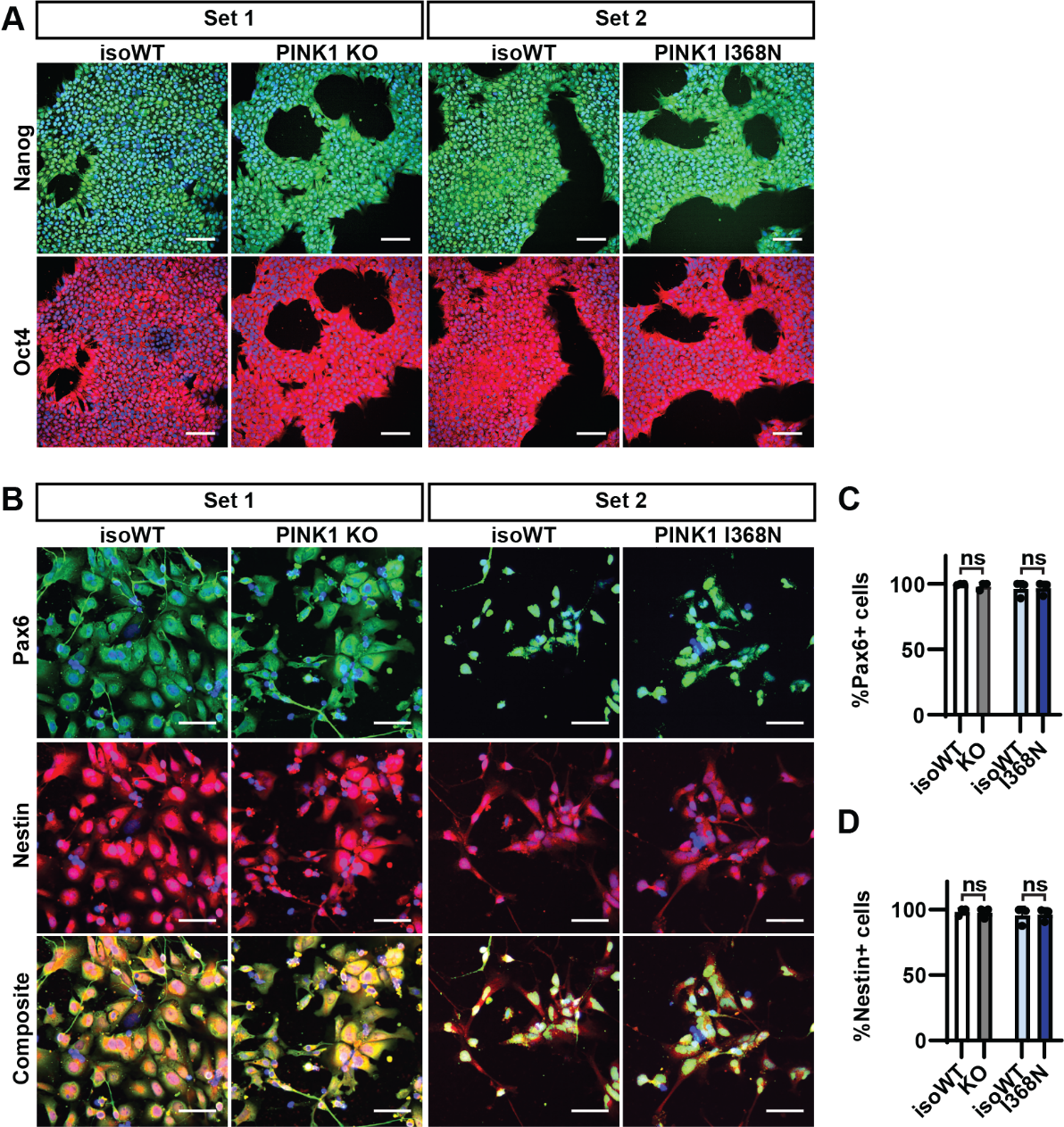

**Fig. S1. iPSC and NPC quality control.** Representative immunofluorescent images showing purity of **(A)** each iPSC set using pluripotency markers Nanog and Oct4. Scale bar = 100 μm and **(B)** each NPC set through quantification of **(C)** Pax6 and **(D)** Nestin. Scale bar = 50 μm. The mean of three independent experiments for each iAS set ± SD is depicted. Data is represented as normalized to isoWT. Data was analyzed by using Student’s T-test. Ns = not significant.

Fig. S2.

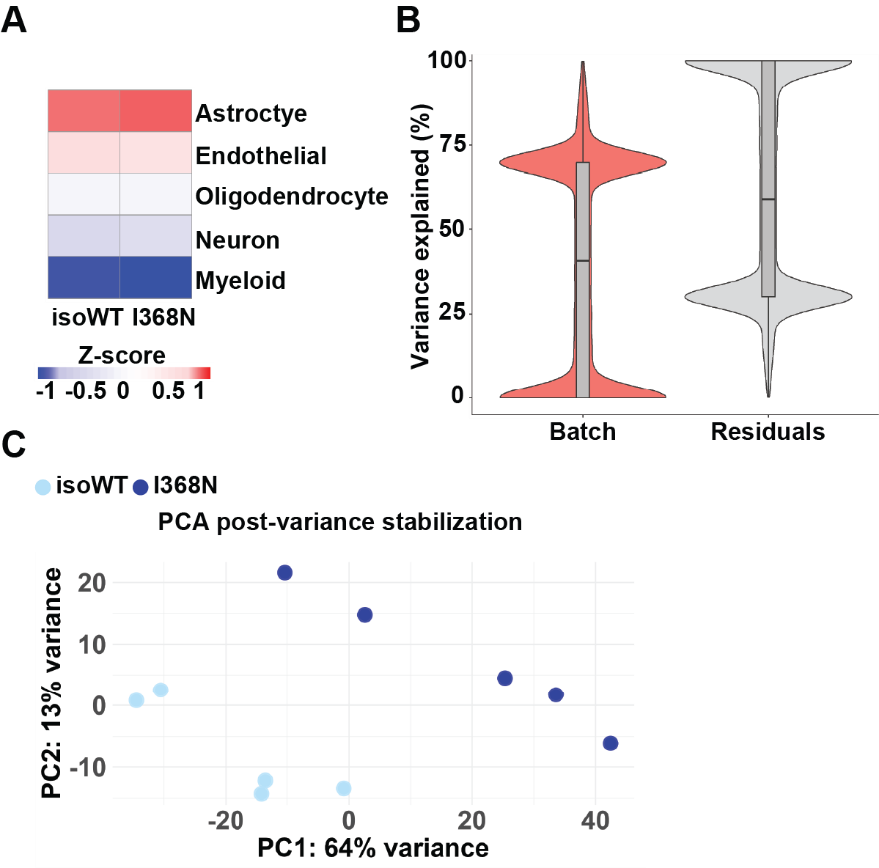

**Fig. S2. Bulk RNA sequencing quality control. (A)** Enrichment analysis of PINK1 I368N and isoWT gene expression compared to various CNS cell types isolated from human brain, using Zhang et al., 2016 dataset. Correlation analysis was performed using Spearman correlation and plotted based on z-scaling. **(B)** Source of variation analysis showing the covariate batch included in the differential expression model and the percentage of gene expression variance it explains. **(C)** Principal component analysis (PCA) was performed on variance-stabilized expression data to assess sample clustering by genotype.

Fig. S3.

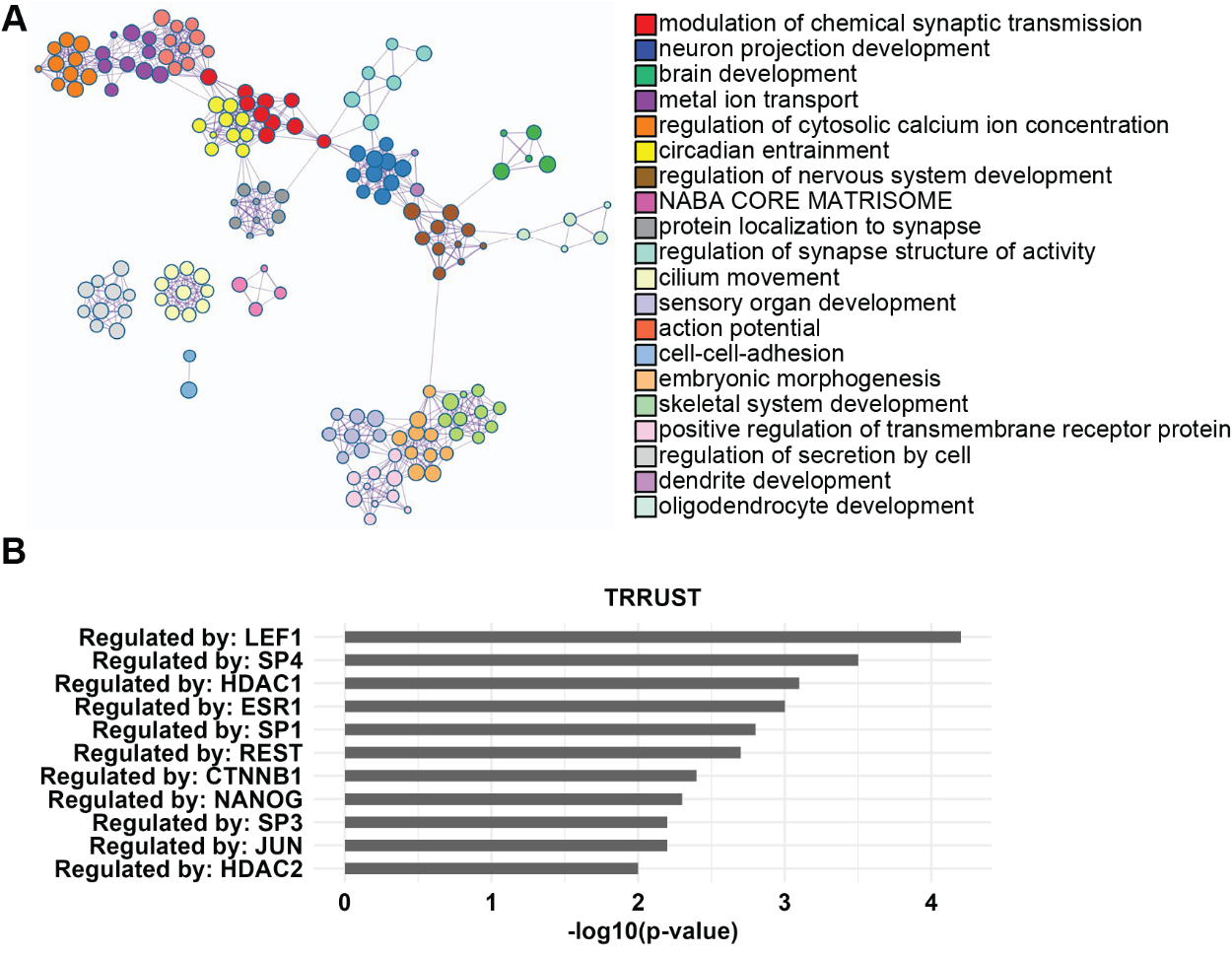

**Fig. S3.** **Independent validation of pathway enrichment and identification of potential upstream transcription factors.** **(A)** Metascape enrichment analysis was performed using all protein coding DEGs, with the top enriched pathways colored by cluster/pathway. **(B)** Transcription factor analysis was performed for all DEGs using the TRRUST database via Metascape.

Fig. S4.

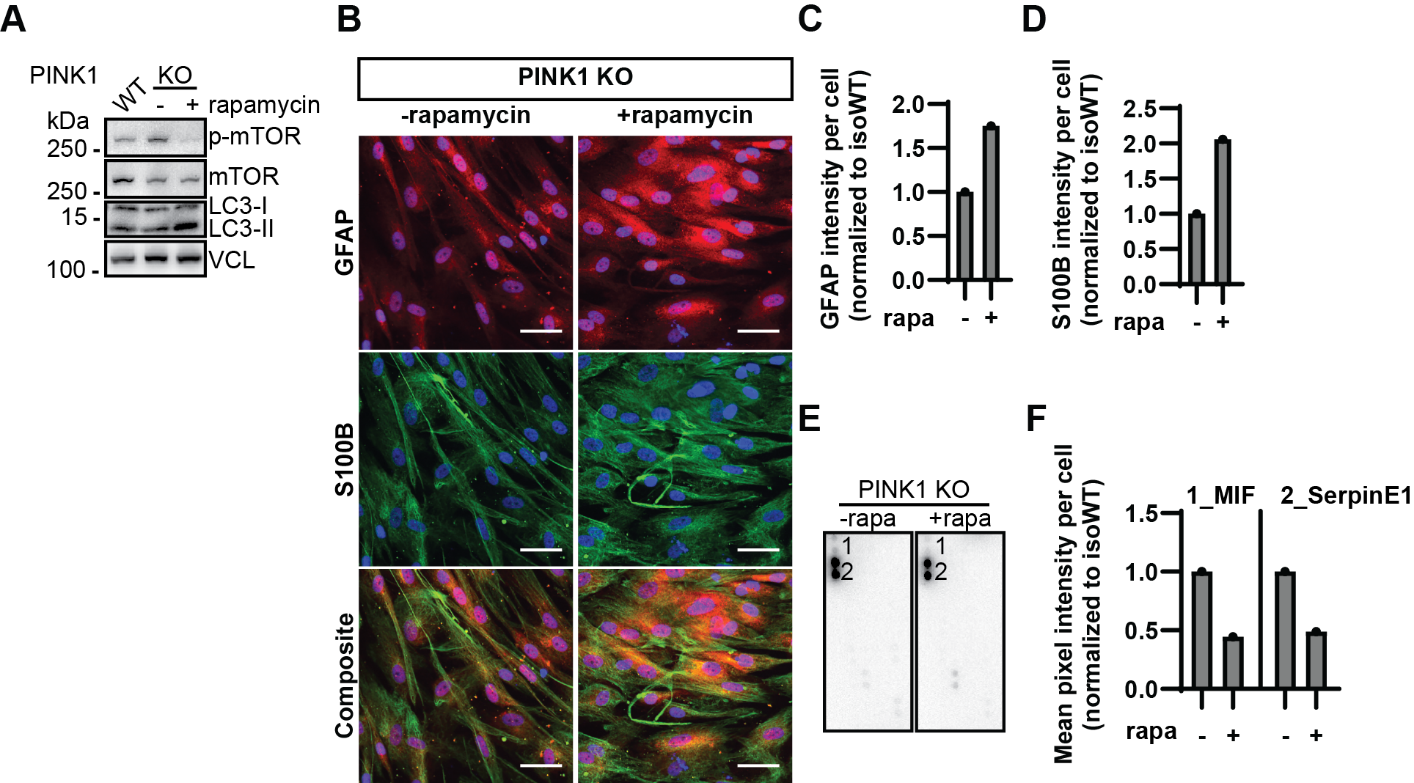

**Fig. S4. PINK1 KO iAS display trend of inflammatory rescue following rapamycin treatment. (A)** Representative western blot from PINK1 KO with and without one week rapamycin treatment and isoWT control showing levels of p-S2448-mTOR, total mTOR, LC3-II, and VCL. **(B)** Immunofluorescence revealed an upregulated trend of **(C)** GFAP and **(D)** S100B in PINK1 KO iAS in response to rapamycin treatment. Scale bar = 50 μm. **(E)** Human cytokine array revealed that rapamycin treatment may rescue **(F)** MIF and SerpinE1 levels in PINK1 KO iAS. The mean of one experiment per assay is depicted, and the data is represented as normalized to PINK1 KO under baseline conditions/no rapamycin treatment.

Fig. S5.

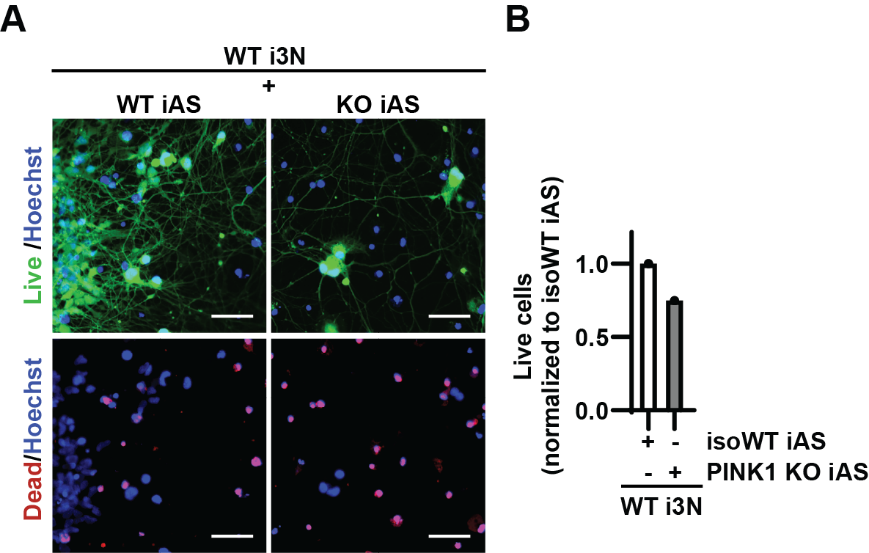

**Fig. S5. PINK1 KO iAS secretome also induces neuronal death.** **(A)** Immunofluorescent imaging of WT i3N cultured with either PINK1 KO or isoWT iAS using LIVE/DEAD assay and **(B)** quantified as the fraction of live cells. Scale bar = 100 μm. Data is represented as normalized to WT i3N cultured with isoWT iAS set to 1. The mean of one experiment is depicted, and the data is represented as the ratio of mean viability.

Table S1.

|  | **Gene symbol** | **Function** | **Enrichment analysis** | **Log2 FC** |
| --- | --- | --- | --- | --- |
| Anti-inflammatory signaling and pathogen recognition | BATF2 | Important regulator of innate immunity. Works to suppress overactive immune signaling from astrocytes^(^*^1-3^*^)^. | GSEA only | -1.210 |
|  | HERC6 | E3 ISGylation ligase that facilitates STING activation. Deficiency attenuates viral recognition^(^*^4^*^)^. | GSEA only | -1.032 |
|  | IFIT3 | Reduced levels previously observed in 1) PINK1 KO primary rodent neurons, 2) PINK1 KO midbrain with A53T synuclein overexpression, 3) human PINK1 KO neuroblastoma cells, 4) PINK1 KO rodent embryonic fibroblasts undergoing acute starvation; upregulated in reactive astrocytes but with a distinct gene expression profile from neurotoxic astrocytes; involved in defense response to viruses^(^*^5-7^*^)^. | GSEA only | -1.063 |
|  | PTGIR | Anti-inflammatory by blocking NF-κB nuclear translocation. Promotes neuroprotection through neuron-astrocyte communication^(^*^8, 9^*^)^. | ORA & GSEA | -2.030 |
|  | TNFRSF1B | Anti-inflammatory TNF receptor. Ablation in mice dysregulates astrocyte-neuron communication^(^*^10-12^*^)^. | ORA & GSEA | -1.202 |
| Proliferation, migration, & cell adhesion | COL13A1^*^ | Plays a role in cell adhesion^(^*^13^*^)^. | ORA only | -2.157 |
|  | NRXN3 | Adhesion molecule that regulates astrocyte morphogenesis and plays a role in cell-cell communication^(^*^14^*^)^. | ORA only | 1.430 |
|  | CCND1 | Downregulation leads to cell cycle arrest in the G0/G1 phase^(^*^15, 16^*^)^. | GSEA only | -1.206 |
|  | CD9 | Common marker for extracellular vesicles; downregulation is associated with increased TNFα levels in astrocytes; also functions as a regulator of proliferation, migration, and cell adhesion^(^*^17-20^*^)^. | GSEA only | -1.071 |
|  | CD70 | Functions as a ligand for CD27 receptor and mediates proliferation in astrocytes^(^*^21^*^)^. | GSEA only | -3.508 |
|  | CD82 | Tetraspanin that also functions to regulate proliferation and migration^(^*^22^*^)^. | GSEA only | -1.368 |
|  | PHLDA2 | Its role has been documented in trophoblasts to inhibit proliferation and migration and induce apoptosis^(^*^23^*^)^. | ORA & GSEA | -1.285 |
| Metabolism | GLRX | Encodes for a cytoplasmic antioxidant enzyme and plays a role in glutathione metabolism^(^*^24^*^)^. | GSEA only | -1.101 |
|  | HK2 | Silencing leads to reduction in p-S65-Ub levels in primary rodent cortical astrocytes. Drives glycolysis; lactate can be released by astrocytes to protect neurons from injury upon stress^(^*^25-27^*^)^. | GSEA only | -1.171 |
| Other | DLG2 | Regulation of hippo signaling pathway and associated with conversion of healthy astrocytes into tumor associated astrocytes in glioblastoma^(^*^28^*^)^. | ORA only | 1.391 |
|  | SNCAIP^*^ | Predicted to interact with amyloid precursor protein and tau. Interacts with alpha-synuclein in neurons^(^*^29^*^)^. | ORA only | 2.801 |
|  | DNAJB4 | Tumor suppressor^(^*^30^*^)^. | GSEA only | -1.026 |
|  | TPBG^*^ | Oncogenic antigen that also regulates dendritic branching and may interact with cytoskeletal proteins^(^*^31^*^)^. | GSEA only | -1.004 |
| Astrocyte-neuron communication | ASIC4 | Voltage-independent proton-gated ion channel. Plays a role in regulating neuron excitability, neurotransmission, and nociception^(^*^32^*^)^. | ORA only | 2.522 |
|  | ATP2B3^*^ | Plays a critical role in Ca^2+^ homeostasis^(^*^33^*^)^. | ORA only | 1.295 |
|  | CHRNB2 | Astrocyte modulation of neurotransmitter release; upregulation is associated with impulsive-like behaviors and cognitive defects in mice^(^*^34^*^)^. | ORA only | 1.274 |
|  | CNGA4^*^ | Voltage-independent cation channel^(^*^35^*^)^. | ORA only | 1.438 |
|  | CNGB1^*^ | Facilitates Ca^2+^ and Na^+^ influx and upregulates neurotransmission^(^*^35^*^)^. | ORA only | 3.880 |
|  | CNR1 | Encodes for the main endocannabinoid effector in the brain; function in mitochondria for regulation of glucose metabolism; upregulation in astrocytes produce time-dependent alterations to lactate levels, which can be mediated through intracellular Ca^2+^ release^(^*^36, 37^*^)^. | ORA only | 1.958 |
|  | CNTNAP2 | Important for neuron–glia interactions; alterations can result in abnormalities in synaptic structure, neuronal network activity, and an autism-related phenotype in mice; upregulation in astrocytes impair glutamate uptake, resulting in neuronal hyperexcitability^(^*^38^*^)^. | ORA only | 2.651 |
|  | ERBB4 | Interacts with NRG3 ligand. The NRG3-ERBB4 signaling pathway elevates astrocyte-neuron interactions in the SNpc of PD patients; upregulated signaling alters cell-cell communication and is implicated in epilepsy; regulator of cell growth; plays a role in cell adhesion^(^*^39-42^*^)^. | ORA only | 2.604 |
|  | GNAO1 | Gain-of-function is associated with movement disorders, while LOF is associated with epilepsy; overexpression lowers Ca^2+^ activity in iPSC-derived astrocytes^(^*^43, 44^*^)^. | ORA only | 2.863 |
|  | GPM6A | Upregulated following brain trauma, such as stroke. Believed to be a protective mechanism to signal synaptic maintenance and membrane remodeling^(^*^45^*^)^. | ORA only | 2.540 |
|  | GRIA2 | Plays an important role in maturation of glutamatergic synapses by replacing Ca^2+^ permeable with Ca^2+^ impermeable AMPA receptors; dysregulation leads to altered synapse number and maturation^(^*^46, 47^*^)^. | ORA only | 3.792 |
|  | GRIA4 | Encodes for an AMPA4 subunit and can mediate astrocyte calcium signaling and astrocyte-neuron interactions^(^*^48^*^)^. | ORA only | 1.502 |
|  | GRIK3 | Ionotropic glutamate receptor subunit that is upregulated in astrocytes in epilepsy^(^*^49^*^)^. | ORA only | 4.005 |
|  | GRIN2A | Encodes for an NMDA receptor subunit^(^*^36, 50^*^)^. | ORA only | 1.964 |
|  | IGSF11 | Regulates synaptic transmission via interactions with PSD95 and AMPA receptors; encodes for a cell adhesion protein; controls gap junction mediated astrocyte-astrocyte communication^(^*^51^*^)^. | ORA only | 2.600 |
|  | KCNC3^*^ | Encodes voltage-gated K^+^ channel for K^+^ buffering and neurotransmission regulation^(^*^52^*^)^. | ORA only | -3.526 |
|  | KCNH2 | Critical for K^+^ homeostasis^(^*^53^*^)^. | ORA only | 1.234 |
|  | KCNJ3 | Plays an important role in Ca^2+^ homeostasis and support for synaptic plasticity^(^*^54^*^)^. | ORA only | 4.257 |
|  | KCNJ13^*^ | Facilitates potassium influx to stabilize cells^(^*^55^*^)^. | ORA only | 5.713 |
|  | KCNMB2 | Induces inhibiting currents via interaction with the large conductance voltage and Ca^2+^-activated K^+^ channel^(^*^56^*^)^. | ORA only | 1.275 |
|  | KCNN3 | Ca^2+^-activated K^+^ channel^(^*^57^*^)^. | ORA only | 1.711 |
|  | LHFPL4^*^ | Tetraspanin that regulates GABA_A_R subunits^(^*^58^*^)^. | ORA only | 2.351 |
|  | LRRTM4 | Overexpression in astrocytes induces synaptic vesicle clustering with axons, leading to postsynaptic currents^(^*^59^*^)^. | ORA only | 1.779 |
|  | P2RX3 | Purinergic receptor activated by ATP; localized to astrocyte processes and may play a role in cell-cell communication^(^*^60^*^)^. | ORA only | 2.767 |
|  | PCDHB13^*^ | Part of the procadherin family, which plays an important role in astrocyte morphogenesis, astrocyte-neuron communication, and synaptogenesis^(^*^61, 62^*^)^. | ORA only | -1.326 |
|  | PRKCG | Becomes activated by Ca^2+^ oscillations, contributing to a negative feedback loop for regulation of neuron excitability; reduces Ca^2+^ spike and wave frequency^(^*^63^*^)^. | ORA only | 1.361 |
|  | SCN2A | Important for astrocyte support for synaptic and ion channel function; downregulated in Alzheimer’s disease patient-derived iAS^(^*^64^*^)^. | ORA only | -1.484 |
|  | SYP | The role in astrocytes is not fully understood, but it could function in synaptic regulation and gliotransmission^(^*^65^*^)^. | ORA only | 1.663 |
|  | GBE1 | Glycogen synthesizer, which is converted into lactate and transmitted to neurons for neurotransmission^(^*^66, 67^*^)^. | GSEA only | -1.085 |
|  | CLCA2^*^ | Regulates Ca^2+^ activated Cl^-^ channels^(^*^68^*^)^. | ORA & GSEA | -2.817 |
|  | EREG | Plays an important role in Ca^2+^ signaling^(^*^69^*^)^. | ORA & GSEA | -2.265 |
|  | RYR2 | Regulator of Ca^2+^ signal transduction and intracellular concentration; increased protein levels or phosphorylation is associated with heightened inflammation^(^*^70, 71^*^)^. | ORA & GSEA | 1.459 |

**Table S1. Functions of leading edge DEGs.** DEGs were defined by adj. p < 0.05 and |Log2 FC| ≥ 1. Pubmed ID’s based on relevance to key word “astrocyte” and most recent publication date were unbiasedly collated using rentrez function via R script, and subsequently summarized and selected based on relevance to astrocytes. ^*^Expressed by astrocytes (Human Protein Atlas), but no publications observed that described its specific role in astrocytes.

**References to Table S1.**

31.	G. R. Jun, J. Chung, J. Mez, R. Barber, G. W. Beecham, D. A. Bennett, J. D. Buxbaum, G. S. Byrd, M. M. Carrasquillo, P. K. Crane, C. Cruchaga, P. De Jager, N. Ertekin-Taner, D. Evans, M. D. Fallin, T. M. Foroud, R. P. Friedland, A. M. Goate, N. R. Graff-Radford, H. Hendrie, K. S. Hall, K. L. Hamilton-Nelson, R. Inzelberg, M. I. Kamboh, J. S. K. Kauwe, W. A. Kukull, B. W. Kunkle, R. Kuwano, E. B. Larson, M. W. Logue, J. J. Manly, E. R. Martin, T. J. Montine, S. Mukherjee, A. Naj, E. M. Reiman, C. Reitz, R. Sherva, P. H. St George-Hyslop, T. Thornton, S. G. Younkin, B. N. Vardarajan, L. S. Wang, J. R. Wendlund, A. R. Winslow, C. Alzheimer's Disease Genetics, J. Haines, R. Mayeux, M. A. Pericak-Vance, G. Schellenberg, K. L. Lunetta, L. A. Farrer, Transethnic genome-wide scan identifies novel Alzheimer's disease loci. *Alzheimers Dement* **13**, 727–738 (2017).
